# FoodAtlas: Automated Knowledge Extraction of Food and Chemicals from Literature

**DOI:** 10.1101/2024.05.16.594596

**Authors:** Jason Youn, Fangzhou Li, Gabriel Simmons, Shanghyeon Kim, Ilias Tagkopoulos

## Abstract

Automated generation of knowledge graphs that accurately capture published information can help with knowledge organization and access, which have the potential to accelerate discovery and innovation. Here, we present an integrated pipeline to construct a large-scale knowledge graph using large language models in an active learning setting. We apply our pipeline to the association of raw food, ingredients, and chemicals, a domain that lacks such knowledge resources. By using an iterative active learning approach of 4,120 manually curated premise-hypothesis pairs as training data for ten consecutive cycles, the entailment model extracted 230,848 food-chemical composition relationships from 155,260 scientific papers, with 106,082 (46.0%) of them never been reported in any published database. To augment the knowledge incorporated in the knowledge graph, we further incorporated information from 5 external databases and ontology sources. We then applied a link prediction model to identify putative food-chemical relationships that were not part of the constructed knowledge graph. Validation of the 443 hypotheses generated by the link prediction model resulted in 355 new food-chemical relationships, while results show that the model score correlates well (R^2^ = 0.70) with the probability of a novel finding. This work demonstrates how automated learning from literature at scale can accelerate discovery and support practical applications through reproducible, evidence-based capture of latent interactions of diverse entities, such as food and chemicals.

## Introduction

Mapping the chemical composition of food and ingredients is essential for unlocking their potential and informing decisions. From creating healthier and tastier food products^1,2^ to enriching food with the right compounds^3,4^ or building personalized diets^5–7^, understanding what is in each ingredient and at what concentration is paramount. Food composition at the molecular level is usually found in food composition tables like the USDA’s FoodData Central (FDC)^8^ or the ANSES-CIQUAL database^9^. This enables several stakeholder groups, from researchers to policymakers, to assess the nutrition quality of various foods and their regulatory status and to use them in the respective industries^10^. However, despite the established importance of the food composition information, most of the food-chemical information that is present in the scientific literature is not captured in the structured databases^1^. For instance, the total size of food composition space is estimated at tens of thousands of chemicals^11^, while FDC and ANSES-CIQUAL focus on only 500 compounds. To expand the coverage of chemicals in foods, several initiatives attempt to capture food composition from scientific literature, such as FooDB^12^ (797 foods and 15,750 detected chemicals) and DietRx^13^ (2,222 foods and 6,992 chemicals), which further aggregate data from several other databases like FDC^8^, KNApSAcK^14^, Dr. Duke’s Phytochemical and Ethnobotanical Databases^15^, Phenol-Explorer^16–18^, and PhytoHub^19^. However, existing databases require laborious annotation effort from experts or lack consistent quality control as the majority of their food-chemical composition information is not linked to evidence that allows reproducible results. For example, less than 1% of associations in FooDB, one of the most notable DBs in this space, have literature citations to support them (**Supplementary Information Section 1.1.1**).

Although manual extraction by the domain experts is often precise, it does not scale well with bibliographic literature sources such as PubMed^20^, which contains 34 million citations and abstracts, and PubMed Central (PMC)^21^, which includes 7.6 million full-text scientific literature articles. From PMC, we estimate we can extract at least 2 million unique food-chemical associations from the unstructured text data (**Supplementary Information Section 1.1.2**). The sheer amount of available scientific literature necessitates the need for an automated framework for constructing knowledge graphs (KGs), which is widely used thanks to their scalability and ability to reveal previously hidden patterns and relationships in the data, leading to better insights and more informed decision-making^22^. There has been prior work utilizing language models to construct domain-specific knowledge graphs from unstructured texts^23–28^, with some combined with active learning (AL) to reduce human annotation^29–31^. Although some works have constructed food-relevant knowledge graphs, they are limited by the low ground truth precision of relation extraction^24^ and the small number of chemical entities^28^.

In this work, we present the Lit2KG framework (**Fig. 1a**) that extracts information from scientific literature using a large language model in an AL setting to construct a largescale KG. The entailment model of the Lit2KG framework uses a premise from the scientific literature to extract and predict multiple hypotheses with high performance (F1 score of 83%), with the predicted probabilities being highly correlated to the ground-truth annotations (R^2^ = 0.94). We also tested four different AL strategies and found that selecting samples that maximize the likelihood leads to discovering new knowledge 38.2% faster than the baseline. Applying graph-embedding link prediction models for graph completion followed by validation through literature search revealed 355 missed food-chemical composition associations that were further verified manually and 11 additional associations that were novel, 6 of which we have found strong evidence to support them. The resulting knowledge graph contains 285,077 triplets of three entity types (food, part, chemical) and four relation types (*contains*, *has part*, *is a*, *has child*) on three evidence quality levels (high, medium, low) with 4,318 of them evaluated by human experts (**Fig. 1b**).

**Fig. 1:**
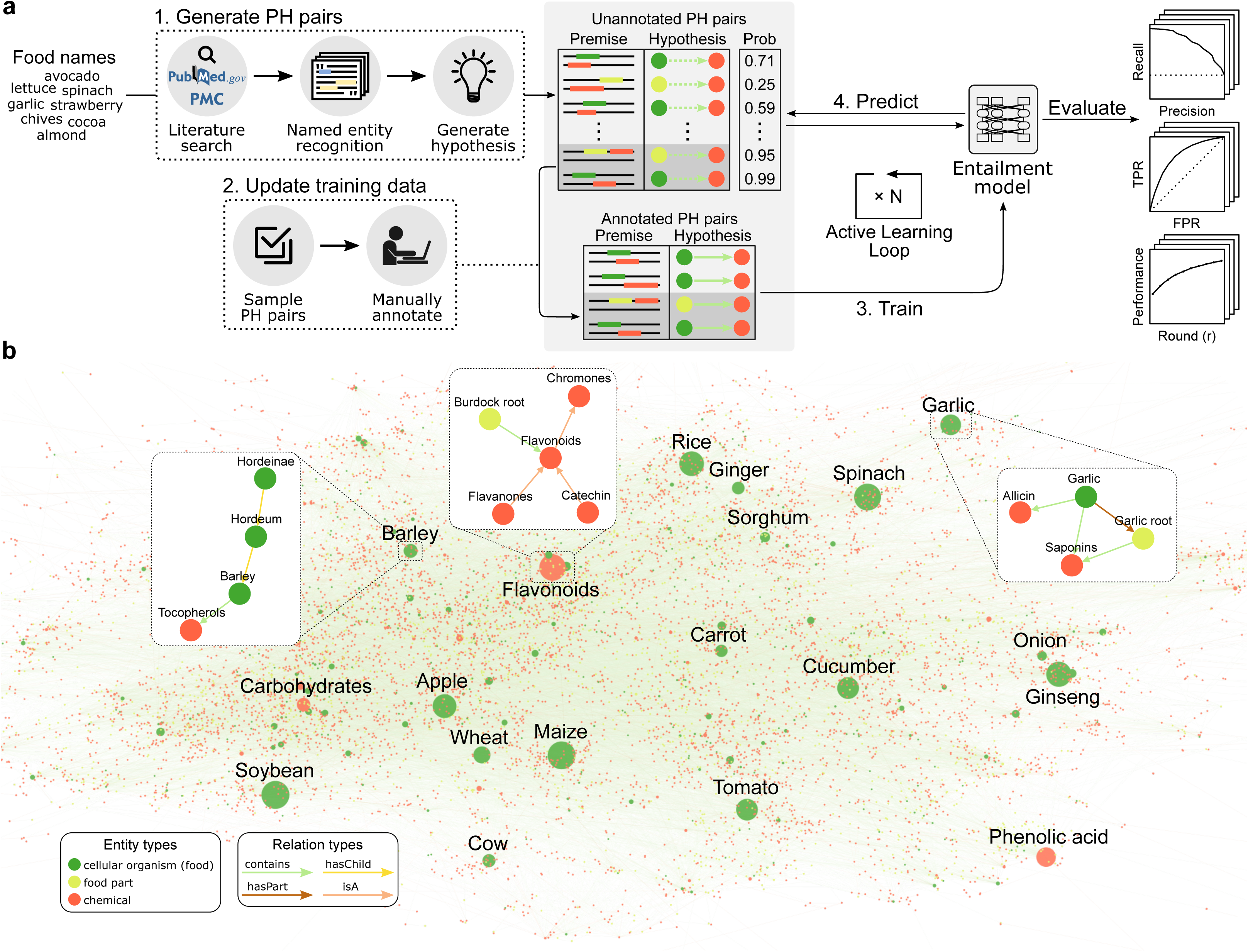
Overview of the Lit2KG framework and the FoodAtlas Knowledge Graph. **a**, Scientific literature is queried using raw food names and retrieved sentences (premises) where the species and chemical entities are tagged (e.g., … cocoa_[SPECIES]_ is a good source of (-)-epicatechin_[CHEMICAL]_ …). From these premises, hypothesis triplets are generated such as (*cocoa, contains, (-)-epicatechin*), which we refer to as premise-hypothesis (PH) pairs. The entailment model is then iteratively updated through active learning cycles, where a new batch of PH pairs is annotated in each cycle. Finally, both annotated and predicted positive PH pairs are used to populate the knowledge graph. **b**, Visualization of the FoodAtlas Knowledge Graph (FAKG), which contains 285,077 triplets of 3 entity types and 4 relation types. Each triplet in the FAKG is assigned one of three quality types and provides a reference to the publications that support it for reproducibility.

## Methods

### Premise-hypothesis pair generation

We collected a total of 1,959 raw and non-processed food names that have a known National Center for Biotechnology Information (NCBI) Taxonomy ID^32^ from multiple food databases. We then used the LitSense API^33^, which is a search system for biomedical literature at the sentence level provided by the NCBI, to query for the search keyword “{*food name*} contains” (**Supplementary Data 1**; **Supplementary Information Section 1.2.1**). The LitSense API returns sentence-level text snippets from the PubMed abstracts and the PMC open-access full-text articles, as well as the named entity recognition (NER) service for species and chemical entities, along with their corresponding NCBI Taxonomy IDs and MeSH IDs, respectively. We further processed these text snippets by discarding non-food entities and tagging the part entities (*e.g.*, leaf and root) using our manually generated lookup table consisting of 70 food parts (**Supplementary Data 2**).

For each LitSense-returned sentence *s*_i_ ∈ *S*, which we refer to as a *premise* in our work, there exist three sets of named entities *F*_i_, *P_i_*, and *C_i_* for food, parts, and chemicals, respectively, where *P*_i_ can be an empty set as not all sentences have parts in them. We then generated a set of hypotheses *H_i_* for each premise *s_i_* by taking the cartesian product of the entity sets *F_i_*, *P_i_*, and *C_i_* as

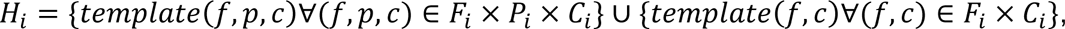

where *template* (⋅) is the hypothesis template that generates a triplet of type ({*food*} {*part*}, contains, {*chemical*}) or ({*food*}, contains, {*chemical*}), respectively. We refer to these pairs of premise and the extracted hypotheses as premise-hypotheses (PH) pairs in our work (see **Supplementary Fig. 1**).

### Premise-hypothesis pair annotation

We annotated the PH pairs to generate a dataset for training, validating, and testing the entailment model using the AL strategy (described in the following sections). During the annotation process, a given PH pair was assigned one of three possible classes *entails*, *does not entail*, and *skip*. More specifically, *entails* was assigned if the premise supported the underlying relationship used to construct the hypothesis, and *does not entail* was assigned if there was insufficient evidence in the premise to support the hypothesis. Note that the hypothesis from a PH pair marked as *does not entail* is not necessarily a negative, as another premise may support the hypothesis. Finally, *skip* was assigned if the premise the LitSense API returned was not formatted correctly or if the NER tagging by LitSense API was wrong (**Supplementary Information Section 1.2.2**). To ensure the annotation was of high quality, two experts annotated each PH pair independently, and only the PH pairs that had agreed annotation results by the two experts were kept. We randomly split the data into training, validation, and test sets with approximate ratios of 70%, 15%, and 15%. To avoid data leakage, we ensured that the three datasets did not share the same premises or hypotheses during the splitting. In the end, we had a training set with 4,120 PH pairs (1,899 *entails*, 2,221 *does not entail*), a validation set with 825 PH pairs (295 *entails*, 530 *does not entail*), and a test set with 840 PH pairs (312 *entails*, 528 *does not entail*) (**Supplementary Data 3**).

### Entailment model

We trained the entailment model to predict whether the premise logically would entail the hypotheses. To this end, we used the BioBERT^34^ over other language models^35–37^ (**Supplementary Information Section 1.2.3**), as it was pre-trained on the same corpus as where the premises were extracted from(PubMed abstracts and PMC full-text articles) and have demonstrated improved performance on biomedical benchmarks^34^. We then fine-tuned the BioBERT entailment model by utilizing the binary classification schema, where the input sequence was formatted by concatenating the premise and hypothesis with the [SEP] token in between, and the model predicted if the given PH pair was *entails* or *does not entail*. We used the held-out validation set to optimize the hyperparameters, where the tunable hyperparameters were learning rate = {2×10^-5^, 5×10^-5^}, epochs = {3, 4}, and batch size = {16, 32}. The hyperparameter set with the best held-out validation precision was selected, and the performance of each round was reported using the held-out test set. Note that we trained a production entailment model using all the labeled data (*i.e.*, training, validation, and test sets) (**Supplementary Information Section 1.2.3**).

### Active learning strategy

In this work, we tested four active learning (AL) strategies, *maximum likelihood*, *maximum entropy*, *stratified*, and *random*. We simulated the AL strategy by splitting the training pool with 4,120 PH pairs into ten rounds *r* = {1, 2, …, 10}, with 412 new PH pairs selected in each round and appended to the existing training data by the respective strategy. In other words, at round *r*, we trained the entailment model *m_r_* using 412 x (*r* – 1) training PH pairs plus 412 new PH pairs selected from the remaining 412 x (10 – *r* + 1) PH pairs. We call this training and evaluation process a *run*, and we repeated 100 *runs* for each AL strategy to test the statistical significance. The *stratified* strategy first ranked the remaining PH pairs from high to low probability and split them into ten equally sized bins, randomly drawing the same number of samples from each bin. The *maximum likelihood* strategy chose the top 412 positive samples based on their probability score. The *maximum entropy* sampling strategy first computed the uncertainty for each PH pair as *min*(1 – *p*, *p*), where *p* is the probability of the given PH pair predicted by the entail model. All PH pairs were then ranked using the uncertainty value from high to low, and the top 412 uncertain PH pairs were selected. Finally, the *random* sampling strategy chose 412 PH pairs randomly. Note that for the first round, all four AL strategies randomly selected the first round of PH pairs to train on, and for the last round, all four AL strategies were trained on a whole training pool of 4,120 PH pairs regardless of the sampling strategy taken. More detailed information can be found in **Supplementary Information Section 1.2.3**, and a visual illustration of the sampling strategies is in **Supplementary Fig. 2**.

**Fig. 2:**
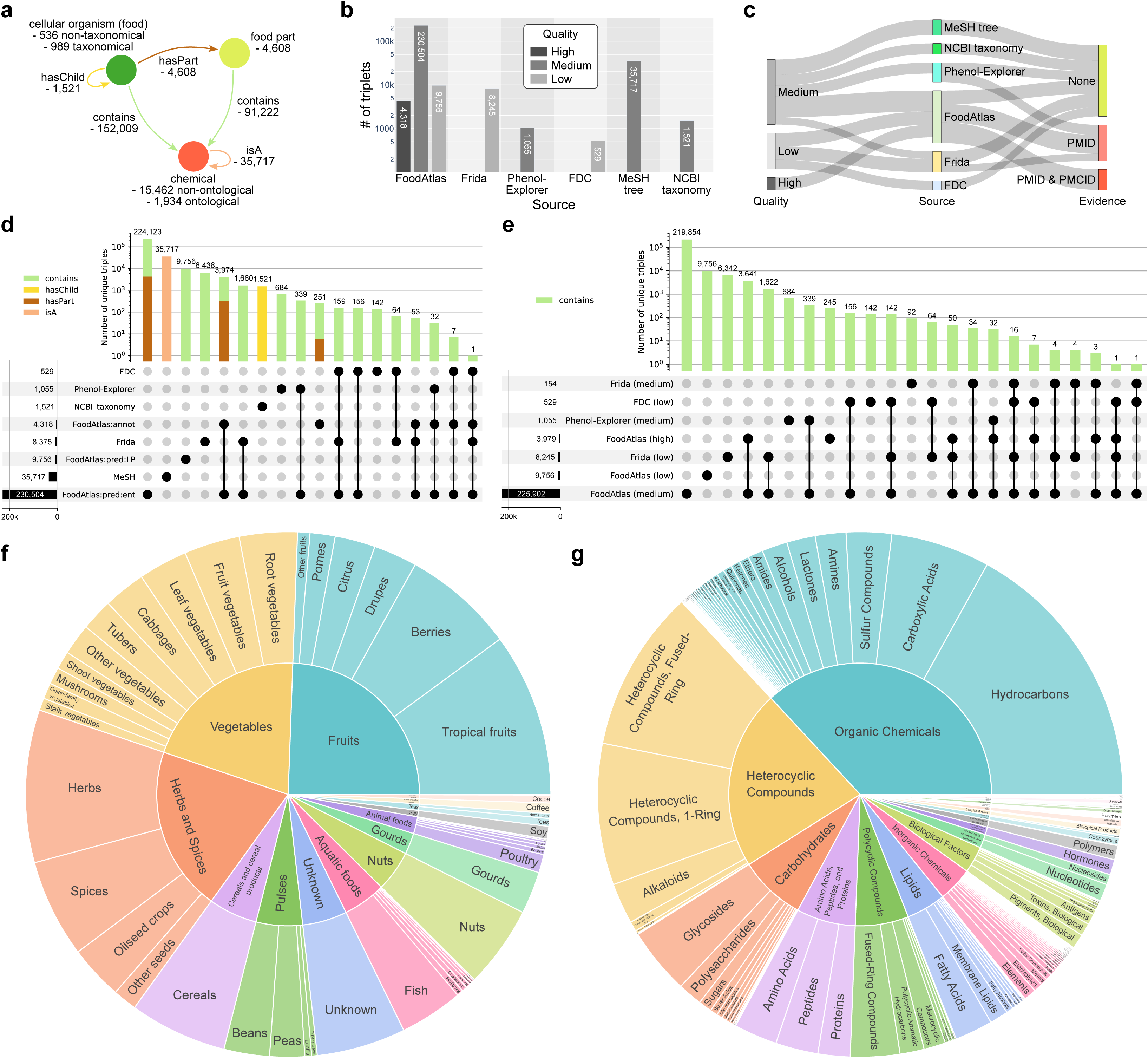
Statistics of the FoodAtlas Knowledge Graph. **a**, Schema of the FAKG. The relation types *contains*, *hasPart*, *isA*, and *hasChild* encode the food-chemical composition relations, the food-food with part relations, the chemical ontological relations using the MeSH tree, and the taxonomical relations using the NCBI Taxonomy, respectively. **b**, Number of triplets per data source in the FAKG depending on the quality. **c**, Sankey graph showing the connections between quality, data source, and evidence. The thickness of the relations between the nodes represents the number of connections in the log scale. **d**, **e**, UpSet plot showing the number of unique triplets for all data sources for all relation types and all sources based on quality for only the contains triplets. Each row in the plot corresponds to a source, and the bar chart on the left shows the size of each source. Each column corresponds to an intersection, where the filled-in cells denote which source is part of an intersection. The bar chart for each column denotes the size of intersections. ‘annot’ stands for annotation, ‘pred’ stands for prediction, and ‘LP’ stands for link prediction. **f**, **g**, Classification of foods and chemicals in FoodAtlas.

### Knowledge graph generation

The FoodAtlas Knowledge Graph *FAKG* = (*E, R*) encodes information using a bag of triplets (*h, r, t*), where {ℎ,*t*} ∈ *E* is the set of all entities (ℎ for the head entity and *t* for the tail entity) and *r* ∈ *R* is the set of all relation. Each triplet in the KG can have one or more sources and qualities. In this work, we define three qualities *high*, *medium*, and *low* for a triplet. The *high*-quality triplets have been validated by the FoodAtlas team and have PMID and/or PMCID. The *medium*-quality triplets are not validated by the FoodAtlas team but have PMID and/or PMCID. Taxonomy and ontology also are medium-quality triplets. The *low*-quality triplets are not validated by the FoodAtlas team and do not have PMID or PMCID. Please refer to **Supplementary Information Section 1.3** for the details of the FAKG design including the entity and relation types.

The first source of information was from the PH pair annotation process, where two relation types, *contains* and *has part*, exist. The triplets with the *contains* relation type were from the positive annotated PH pairs, whereas the triplets with the *has part* relation type were automatically extracted from the *contains* triplets. For example, a triplet (*coconut, has part, coconut seed*) was extracted from the triplet (*coconut seed, contains, lauric acid*). All triplets from this source were high-quality. The second source was the entailment model predictions, also with the *contains* and *has part* relation types. However, these were not annotated and thus were assigned a medium-quality. The third source was the enrichment through the NCBI Taxonomy and MeSH tree ontology. The NCBI Taxonomy, which contains medium-quality triplets with the *has child* relation type, encodes the hierarchical structure of the taxonomic lineage (*Cocos (genus), has child, Cocos nucifera (species)*). The MeSH tree, which contains medium-quality triplets with the *is a* relation type, encodes the ontological relationship of the chemical entities. We also included the triplets extracted from the external databases (Frida^38^, FDC, and Phenol-Explorer) with either *medium*- or *low*-quality triplets with the *contains* relation type. Finally, we also included the link prediction results (triplets with the *contains* relation type) as low-quality.

### Link prediction

Link prediction is a widely studied field that refers to the task of predicting missing relationships or links between entities in a graph, (food, contains, chemical) triplet type in our case, and contributes to the enhancement and enrichment of knowledge graphs^39^. Using the Python library PyKEEN^40^, we trained a set of benchmark link prediction models TransE^41^, ER-MLP^42^, DistMult^43^, TransD^44^, ComplEx^45^, and RotatE^46^ on different versions of the FAKG (**Fig. 5a,b**), performed hyperparameter optimization on the held-out validation set using mean rank (MR), and reported the results on the held-out test set (**Supplementary Information Section 1.2.4**). The link prediction models were also calibrated using isotonic regression to provide an interpretable probability score. Link prediction models are commonly evaluated using rank-based metrics like mean rank (MR), mean reciprocal rank (MRR), hits@1, hits@3, and hits@10^47^. However, our end goal was to generate hypotheses that were either true or false, and therefore, we decided to also evaluate using standard binary classification metrics like confusion matrix, precision, and recall. To this end, we randomly sampled two negatives for each positive triplet in the validation and test set by corrupting the head and tail entity once, which resulted in a validation set with 1,335 triplets (445 positives and 890 negatives) and a test set with 1,341 triplets (447 positives and 894 negatives). Due to the nature of the graph-embedding models that cannot make predictions on test triplets with an entity that is never seen during the training, we report our binary classification metrics in a stricter *unfiltered* setting, where the test triplets that would be dropped in the *filtered* setting are kept and assigned a default majority label 0.

### Link prediction literature validation

To validate the link prediction-generated food-chemical triplets, we searched the following four sources sequentially: PubChem taxonomy^48^, Bing Chat, Google Scholar, and Google. Specifically, for a given food-chemical pair, we first checked if the Taxonomy section of PubChem entry for the chemical of interest lists the scientific name of the food and has a reference. If not, we then asked Bing Chat, a search engine based on a large language model, to find the reference (**Supplementary Fig. 3**). Next, we searched Google Scholar using a set of pre-defined search queries (**Supplementary Information Section 1.1.8.2**). If the initial Google Scholar search did not return the positive relationship within the first three pages (30 papers, 10 papers per page), we repeated the process with the synonyms of the entities. Finally, we searched the first 30 contents of Google using the same search method as Google Scholar. A complete procedure for the link prediction validation can be found in **Supplementary Information Section 1.1.8**.

**Fig. 3:**
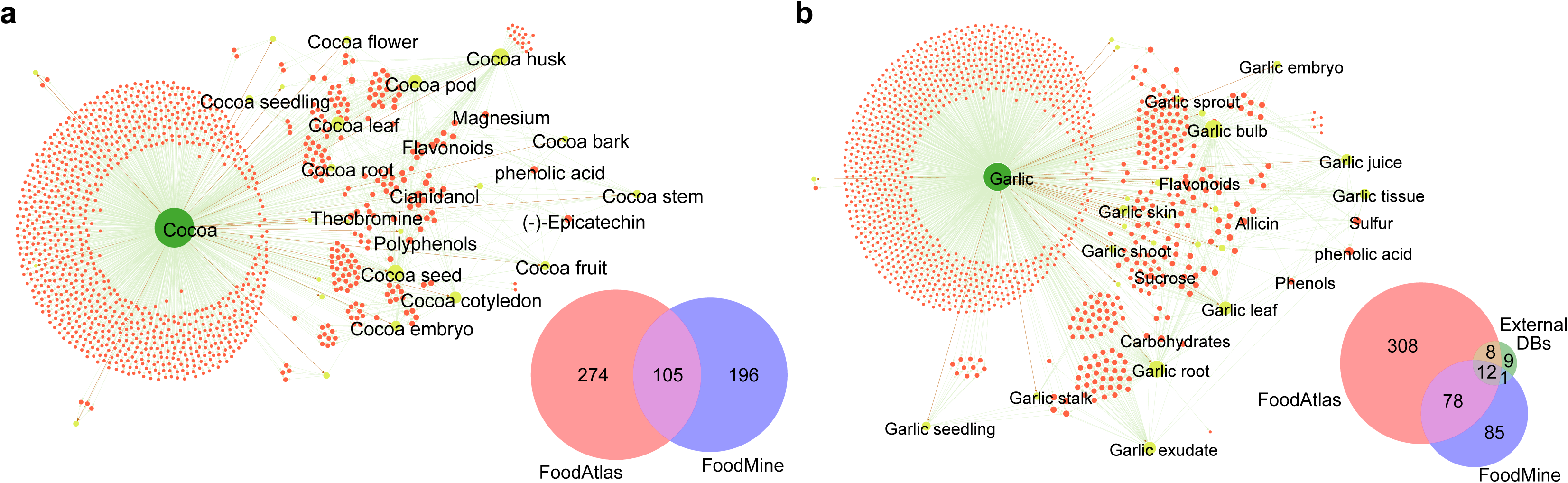
Results of comparing cocoa and garlic to the benchmark dataset FoodMine. **a**, **b**, FoodAtlas subgraph of cocoa and garlic where whole food and food parts and their chemical composition are displayed. The label of the top 20 nodes with the largest degree is shown for each subgraph, and the size of the node is proportionate to its degree. The Venn diagram shows the overlap of FoodAtlas (entailment model annotation, entailment model prediction, and link prediction), external databases (Frida, Phenol-Explorer, and FDC), and FoodMine. Interestingly, none of the 3 external databases reported any chemical composition of cocoa.

## Results

### The FoodAtlas Knowledge Graph contains a wide spectrum of food-chemical composition information

We utilized the Lit2KG framework (**Fig. 1a**) to extract the food-chemical composition information from the PubMed abstracts and open-access articles using raw food ingredients as queries (see **Methods**). From this search, we generated 3,596,755 premise-hypotheses (PH) pairs where the hypotheses are (food, contains, chemical) or (food part, contains, chemical) triplets. We then used BioBERT^34^, a biomedical language representation model for triplet binary classification that we fine-tuned with 4,318 manually curated positive triplets in an active learning setting. This resulted in 230,504 additional positive triplets, for a total of 234,822 unique positive triplets. In addition, we curated and added the food-chemical composition information based on quality criteria from three external databases (8,375 triplets from Frida^38^, 1,055 triplets from Phenol-Explorer^18^, and 529 triplets from FDC^8^), taxonomical information of the foods using the NCBI Taxonomy (1,526 triplets), and ontological information of the chemicals using the MeSH tree (43,691 triplets) (**Fig. 2d**). Applying link prediction on the knowledge graph generated an additional 9,756 triplets of food and chemical pairs, 355 of them manually validated as positives. The final FoodAtlas knowledge graph (FAKG, **Fig. 1b**) contains 536 food entities, 4,608 food parts, 15,462 chemical entities, and 285,077 unique triplets about food-chemical composition with four different relation types and three different entity types (**Fig. 2a-g**).

In terms of triplet quality, FAKG has 4,318 (1.5%) high-quality (*i.e.*, validated by two experts), 264,455 (92.8%) medium-quality (*i.e.*, with at least one reference, but not manually validated), and 16,304 low-quality (5.7%) triplets (*i.e.*, no references, see **Methods** and **Fig. 2b**). From those, 4,318, 226,437, and 9,756, respectively, have been uniquely captured by our Lit2KG pipeline and the link prediction analysis (**Supplementary Information Section 1.1.3** and **Fig. 2b,c**). The top five foods whose chemical composition is most well documented in the knowledge graph are soybean (Glycine max), maize (*Zea mays*), rice (*Oryza sativa*), cucumber (*Cucumis sativus*), followed by tomato (*Solanum lycopersicum*) (**Supplementary Fig. 4**).

### FoodAtlas discovers complementary information to benchmark datasets

To test how good the coverage of the food-chemical composition triplets from the Lit2KG pipeline is, we compared them with FoodMine^49^, a database that contains a manually curated chemical composition of two selected foods, cocoa (592 chemicals) and garlic (289 chemicals). Although there were initially 1,289 cocoa and 1,376 garlic chemicals in FAKG, we adopted the same method used by FoodMine to make chemicals in the two sources comparable and created an additional chemical identifier, specifically for matching FoodMine chemicals with those in FAKG (**Supplementary Information Section 1.1.4**). After this processing step, the FAKG has 379 cocoa and 406 garlic chemicals, whereas FoodMine has 301 and 176, respectively. Out of 575 chemicals for cocoa, 274 (47.7%) chemicals were found in FAKG but not in FoodMine, 105 (18.3%) chemicals were common between the two, and 196 (34.1%) chemicals were not found in FoodAtlas (**Fig. 3a**). For garlic, FoodAtlas was able to capture 51.1% (90 out of 176) of FoodMine chemicals, while 316 chemicals were unique to FAKG (**Fig. 3b**; see **Supplementary Fig. 5** for a similar comparison with FooDB).

### Maximum likelihood active learning strategy discovers knowledge 38% faster than without

We fine-tuned the BioBERT-based entailment models based on four different AL strategies over ten rounds (see **Methods**; **Supplementary Fig. 6**). Although all four AL strategies eventually discovered the same set of 1,899 positives among the 4,120 PH pairs in the training pool at the final round (*r* = 10), the maximum likelihood strategy identified the positives in training set by 38.2% ± 27.3% faster than the active learning baseline of choosing random pairs, followed by the maximum entropy (10.7% ± 6.6%) and stratified learning (9.3% ± 5.3%; **Fig. 4a,b** and **Supplementary Information Section 1.1.5**). Concomitantly, we observed lower performance for the entailment models trained using the maximum likelihood strategy than the others on all metrics for rounds 2 through 4 (adjusted *p*-value < 3.6 × 10^-2^). This was due to data imbalance, as the maximum likelihood strategy samples PH pairs that were highly probable, and thus its entailment models were trained on an unbalanced training set where on average, 74.9% of the training data for rounds 2 through 4 was positive compared to 53.6%, 52.2%, and 46.1% for maximum entropy, stratified, and random, respectively, (**Supplementary Fig. 7**).

**Fig. 4:**
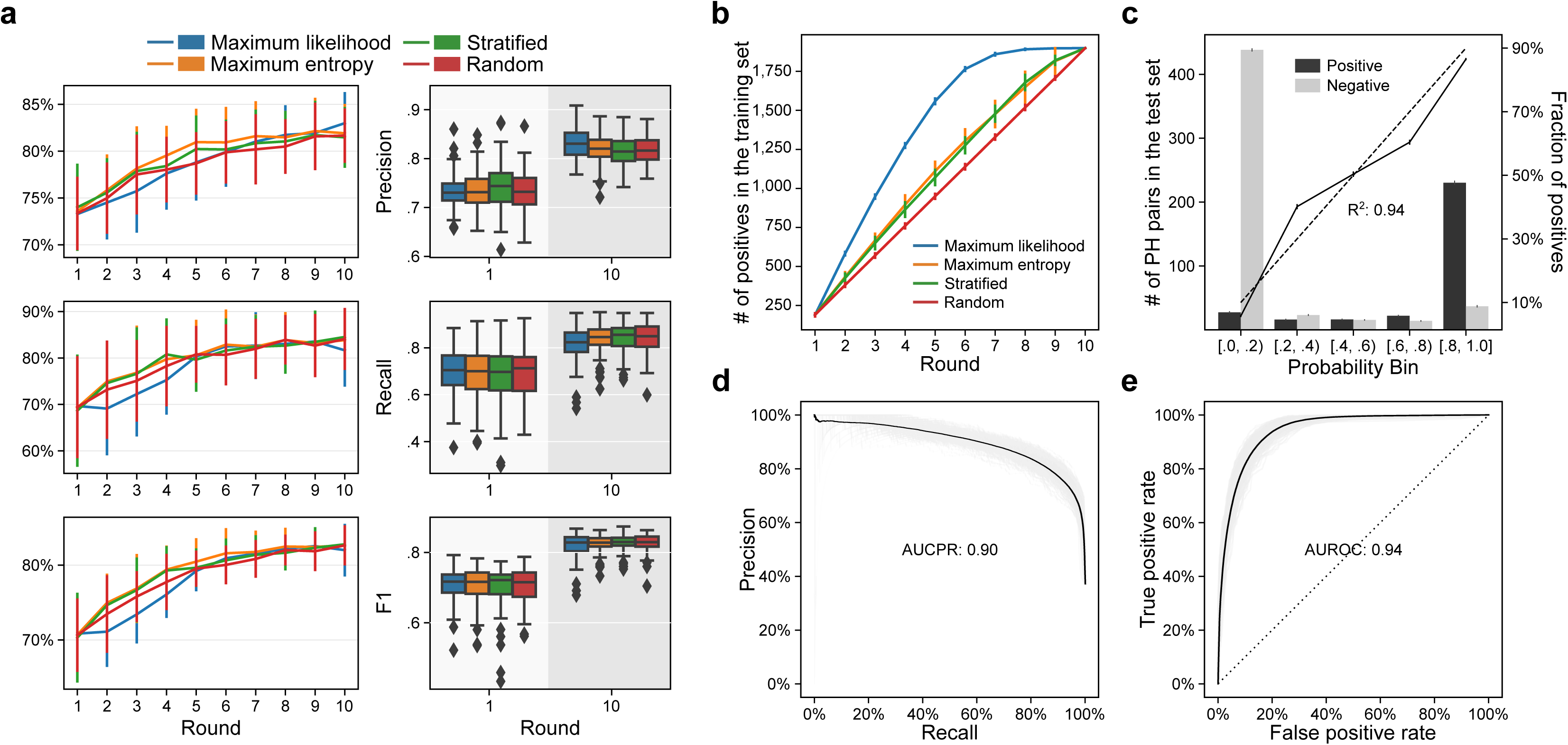
Prediction performance of the entailment model. **a**, Precision, recall, and F1 score of the entailment models trained using the 4 different AL strategies for initial (r = 1) and final (r = 10) rounds (n = 100, 100 different random seeds). On the left, the line plot shows the mean value of each AL strategy, and the error lines denote the standard deviation of the 100 random seeds. On the right, the box represents the interquartile range, the middle line represents the median, the whisker line extends from minimum to maximum values, and the diamond represents outliers. **b**, Comparison of the new knowledge discovery rate compared between the 4 AL strategies. The plot shows how early on in the AL round the 1,899 positive triplets within the simulated training pool of 4,120 triplets are discovered. The error line shows the standard deviation of the 100 random seeds. **c**, Calibration plot showing a high correlation between the probability assigned by the entailment model and the ground-truth annotations on the test set (R^2^ = 0.94). **d**, **e**, The precision-recall and receiver operating characteristic curves of the entailment model predictions compared to the ground-truth annotations in the test set at the final round (r = 10) averaged over all 400 runs with a different random seed (100 runs for each of the 4 AL strategies).

For the final entailment models, AUCPR = 0.90 and AUROC = 0.94, where the baselines were 0.37 and 0.50, respectively (**Fig. 4d,e** and **Supplementary Table 1**). The model PH prediction probability was well-calibrated and highly correlated with the actual ground truth statistics after manual validation (R^2^ = 0.94, **Fig. 4c**). For instance, 88.6% out of all triplets with a probability ≥ 0.9 were positives, whereas only 3.9% with a probability < 0.1 were positives (**Supplementary Fig. 8**).

### Sources of error and impact of large language model general knowledge

Not surprisingly, the entailment model predicted best on straightforward, simple sentence structure, while its performance deteriorated when domain expertise was needed or premises were hypotheses posed by the authors as shown in index 5-8 of **Table 1** (see **Supplementary Information Section 1.1.6**). Variance across bootstrapped models was maximized with uncertainty: predictions with 40% to 60% probability had a standard deviation of 0.31 vs. 0.04 for predictions with less than 10% or more than 90% probability (*p*-value = 2.2 x 10^-177^; see **Supplementary Data 3**). Furthermore, analyzing the entailment model prediction results based on which section of the literature the premise was taken from (*e.g.*, introduction, methods, etc.) revealed higher precision in certain sections. Unexpectedly, hypotheses stemming from the introduction and methods sections were associated with high precision (0.91 and 0.89, respectively) when compared to sections like abstract, title, and conclusion (0.77, 0.75, and 0.74, respectively; *p*-value: 9.7 x 10^-76^) (see **Supplementary Information Section 1.1.7** and **Supplementary Table 2**).

**Table 1.** Comparison of the entailment model predicted premise-hypotheses pairs and the ground-truth annotation. The probability column shows the mean and standard deviation of the probability scores assigned to the corresponding PH pair at the final round (r = 10) of active learning by the 400 entailment models (100 random seeds each for 4 active learning strategies). GT stands for ground truth class assigned by the consensus of two annotators based on the premise. Samples shown in this table are from the test set.

| Index | Premise | Hypothesis | Section | GT | Prediction | Probability |
| --- | --- | --- | --- | --- | --- | --- |
| 1 | Standardized extracts from the **leaves** of **Ginkgo biloba** contains 24 % ginkgo-flavone glycosides and 6 % terpenoids (**ginkgolides**, bilobalide).<sup>77</sup> | (Ginkgo biloba – leaves, contains, ginkgolides) | Intro | Entails | Entails | 99.6% ± 1.0 % |
| 2 | This Vaccinium myrtillus L extract is composed of **flavonoids**, and standardized to contain 36% anthocyanins, with conformance to the USP 31 on 'Powdered **Bilberry** Extract'.<sup>78</sup> | (Bilberry, contains, flavonoids) | Methods | Entails | Entails | 51.3% ± 33.7% |
| 3 | RYNXC consisted of 9 traditional Chinese herbs, including **clove**, rhubarb, frankincense, myrrh, **borneol**, rhizoma corydalis, cowherb **seed**, Rosae rugosae, Garden balsam stem.<sup>79</sup> | (Clove – seed, contains, borneol) | Intro | Does not entail | Does not entail | 0.3% ± 0.4% |
| 4 | For this purpose, tablets were produced containing 16 mg of **ellagic acid** with 100 mg of pulp from the fruit of an evergreen tree called Cherimoya, sour-sop, **custard apple**, and other common names (Annona muricata).<sup>80</sup> | (Custard apple, contains, ellagic acid) | N/A | Does not entail | Does not entail | 44.4% ± 37.0% |
| 5 | Previous investigations postulated that **polyunsaturated fatty acids** (PUFAs) are essential nutrients for the **common octopus**.<sup>81</sup> | (Common octopus, contains, polyunsaturated fatty acids) | Intro | Does not entail | Entails | 99.3% ± 1.0% |
| 6 | **Domoic acid** excretion in **dungeness crabs**, razor clams and mussels.<sup>82</sup> | (Dungeness crabs, contains, Domoic acid) | Title | Does not entail | Entails | 62.7% ± 34.4% |
| 7 | Antihyperlipidaemic and antihypercholesterolaemic effects of **Anethum graveolens** leaves after the removal of **furocoumarins**.<sup>83</sup> | (Anethum graveolens, contains, furocoumarins) | Title | Entails | Does not entail | 2.3% ± 8.0% |
| 8 | In the study by Keskiner et al. (2017), the patients in the test group received capsules containing 6.25 mg EPA and 19.19 mg **DHA** from **Atlantic salmon** (Vectomega tablet, Laboratoires Le Stum, Plage, France).<sup>84</sup> | (Atlantic salmon, contains, DHA) | Results | Entails | Does not entail | 44.7% ± 34.3% |

### Link prediction, GPT model, and the impact of ontologies in performance

We trained a set of link prediction models for the *contains* relation between previously unknown food-chemical pairs (**Fig. 5a**). The best performance was from TransD trained on the FA_A,R_ (TransD-FA_A,R_) with an overall best performance (precision: 79.3%, recall: 75.4%, and F1: 77.2%) (**Fig. 5b**). However, as these models cannot classify triplets with entities not seen during the training phase, we used the next best model, RotatEFA_A,E,R,P80_ that has this capacity (precision: 76.8%, recall: 70.6%, and F1: 73.5%; **Supplementary Data 4** and **Fig. 5c**). Interestingly, the inclusion of ontological information (Enrichment in **Fig. 5b**), increases the F1 score by 22.2% (63.2% of FA_A_ vs. 77.2% of FA_A,R_; *p*-value = 2.4×10^-5^). Moreover, RotatE-FA_A,E,R,P80_ is highly calibrated with R^2^ = 0.99 (**Fig. 5d**) and has an AUCPR of 0.82 (baseline 0.33) and AUROC of 0.88 (baseline 0.5) (**Fig. 5e,f**). All link prediction models performed better than the generalized GPT-3.5 model (text-davinci-003), which was not fine-tuned using the KG (precision: 64.8%, recall: 31.8%, and F1: 42.7%) (**Supplementary Information Section 1.2.4**).

### Link prediction reveals previously unknown food-chemical relationships

The final FAKG contains 536 food entities (excluding food part entities) and 15,462 chemical entities, which translates to 8,287,632 possible food-chemical pairs. Only 1.72% (142,253 triplets) of these food-chemical pairs are connected via the *contains* relation, with the rest, 98.28% (8,145,379 triplets), being unknown. We, therefore, used RotatE to assign probability scores to these unknown pairs (**Fig. 6a**), among which 9,756 pairs (0.1%) were assigned a positive prediction label (see **Methods** and **Supplementary Data 5**). Validating 443 sampled hypotheses from these pairs through an extensive literature search (**Fig. 6b** and **Supplementary Information Section 1.1.8**) revealed 355 positive *contains* triplets between 203 foods and 153 chemicals (**Fig. 6c**), while 11 triplets remained yet unknown with no direct evidence (**Supplementary Table 3** and **Supplementary Data 5**).

**Fig. 5:**
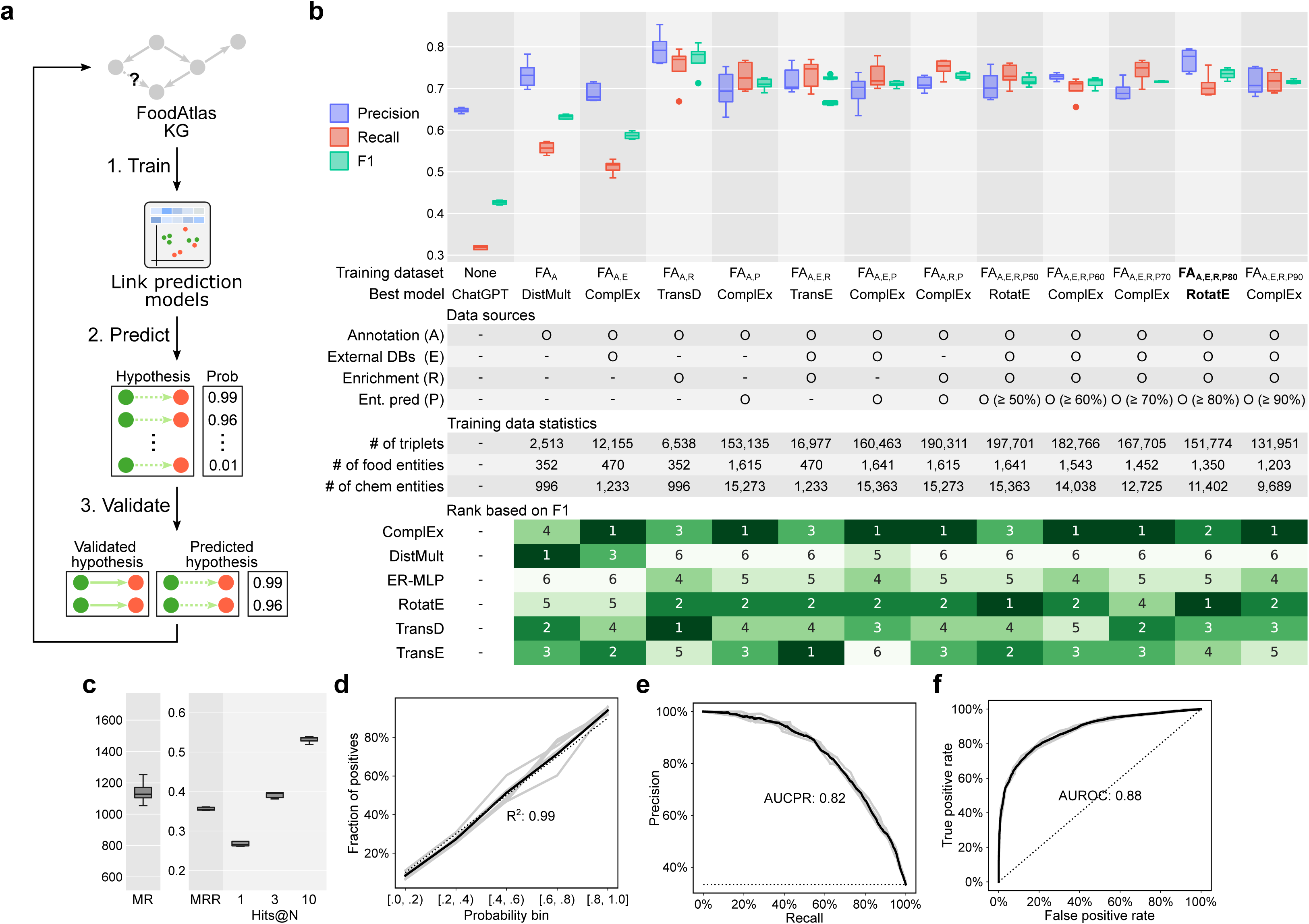
Link prediction model performance. **a**, We use the FAKG to train a link prediction model whose objective is to generate hypotheses of type (food, contains, chemical) that is previously unknown in the graph. **b**, Ablation study result showing the performance of 6 different link prediction models trained using 12 different versions of the FAKG, where different data sources were added or removed to understand their importance. While the training data is different for each version of the dataset, the validation and test set remain the same for fair comparison (positive to negative ratio is 1 to 2; baseline precision: 0.33, recall: 1.0, F1: 0.46). The best model for each dataset is selected based on the F1 score. The box represents the interquartile range, the middle line represents the median, the whisker line extends from minimum to maximum values, and the diamond represents outliers. **c**, Standard rank-based metrics of the best model (RotatE) trained on the best training dataset (FA_A,E,R,P80_). Lower is better for mean rank (MR), while higher is better for mean reciprocal rank (MRR), hits@1, hits@3, and hits@10. **d**, Calibration plot showing a high correlation between the probability assigned by the link prediction model and the ground-truth annotations on the test set (n = 5, 5 different random seeds). **e**, **f**, Precision-recall and receiver operating characteristic curves of the best link prediction model.

**Fig. 6:**
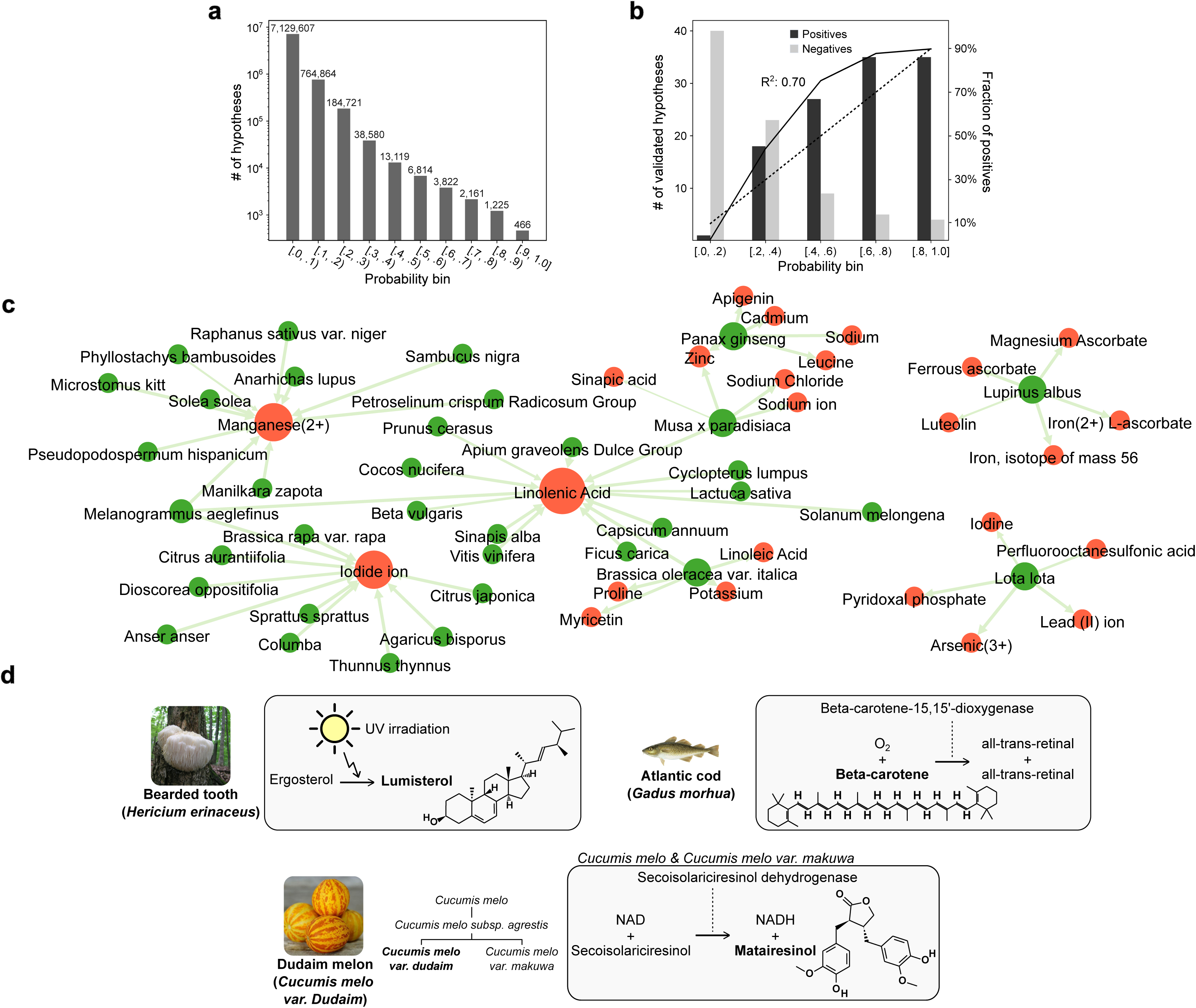
Validation of link prediction generated hypotheses. **a**, Distribution of the 8,145,379 hypotheses in 10 equally spaced bins. **b**, Calibration plot of the link prediction model based on randomly selected hypotheses (40 per bin) validated through manual literature search. **c**, Visualization of positively validated link prediction hypotheses, where the 1-hop subgraph of the top 3 chemical and 5 food entities are shown. The edge width is proportionate to its probability score, and the size of the node is proportionate to its degree. **d**, Indirect evidence for the 6 food-chemical relationships not found in the manual literature search and suggested by the link prediction pipeline, where the food and chemical of interest are marked in bold.

A closer look at the 355 triplets demonstrated the importance of link prediction for knowledge graph completion. Linolenic acid, which is an essential omega-3 fatty acid that must be obtained through the diet and helps reduce inflammation^50^, lower blood pressure^51^, and improve cholesterol levels^51^, were validated to be found in 14 different foods (**Fig. 6c**). The link prediction also discovered evident relationships such as the iodide ion, which is an essential trace element for vertebrates, and manganese(2+), which is a cofactor for many enzymes involved in metabolism^52^, including those that are important for bone development^53^ and antioxidant defense^54^, each with relationship to 10 different foods (**Fig. 6c**). When it comes to foods, we identified five foods, *Lota lota* (NCBI:txid69944), *Brassica oleracea var. italica* (NCBI:txid36774), *Lupinus albus* (NCBI:txid3870), *Panax ginseng* (NCBI:txid4054), *Musa x paradisiaca* (NCBI:txid89151), that have largest number of positively validated positives to 5 chemicals each (**Fig. 6c**).

### AI-driven discovery of six food-chemical relationships

We performed additional analysis for the 11 potential novel food-chemical candidates not reported in the literature (**Supplementary Table 3**) and found strong evidence that supports the relationships for 6 of them. **Fig. 6d** shows 3 of these potentially novel food-chemical relationships, whereas the rest can be found in **Supplementary Fig. 12** and **Supplementary Information Section 1.1.8**. For (Atlantic cod, beta-carotene), metabolic pathway analysis identified homologous enzymes directly associated with the synthesis or metabolism of the chemical in the food (**Supplementary Information Section 1.1.8**). Specifically, the enzyme beta-carotene-15,15’-dioxygenase, which metabolizes beta-carotene in human^55^, had 58.5% sequence similarity with beta,beta-carotene 15,15’-dioxygenase-like in the Atlantic cod. Similarly, for (dudaim melon, matairesinol), we found an enzyme secoisolariciresinol dehydrogenase for biosynthesis of matairesinol in genetically close species *Cucumis melo* and varietas *Cucumis melo var. makuwa*^56^, as we did not have the *Cucumis melo var. dudaim* genome to run a direct search. For the (bearded tooth, lumisterol) pair, we found the existence of ergosterol in bearded tooth^57^ that converts to lumisterol under UV irradiation^58^.

## Discussion

In this work, we created an automated framework to extract information from literature and create domain-specific knowledgebase graphs. Applying to food and chemical relationships created the first AI-driven resource in the field, summarizing findings through 285,077 triplets, with 106,082 (2,091 high-, 94,095 medium-, and 9,896 low-quality) of those associations (46.0%) never been reported before in published databases (**Supplementary Information Section 1.1.9**). While 98.2% of triplets from the Lit2KG pipeline were labeled as either medium or low quality (**Fig. 2b**), our results indicate high performance for both the entailment model (medium-quality triplets; precision of 0.82) and the link prediction model (low-quality triplets; precision of 0.77). Additionally, both models exhibit strong calibration (R^2^ of 0.94 and 0.99, respectively); that is, the model’s predicted probabilities accurately reflect the likelihood of outcomes, providing reliability, interpretability, and better decision support. Surprisingly, in many cases, there are no indexed references associated with the reported entries and unique standardized IDs for the foods and compounds, which made reproducibility and provenance very difficult (**Supplementary Information Section 1.3.3** and **Supplementary Table 6**). FoodAtlas, by design, addresses this challenge by associating one or more references to each association.

Similarly, we are surprised that most of the associations that we have mined from the literature are not part of the existing databases, which argues that there is a plethora of information to be identified, validated, and integrated into tools like FoodAtlas. This, in turn, will be a boon for data-driven tools and pipelines for various applications, compound and source identification, product formulations, and other R&D operations that currently are serendipitous, error-prone, and time-consuming. Concomitantly, the food-chemical composition knowledge coverage of what is currently in various databases varies (22% of Frida, 35% of Phenol-Explorer, 61% of FDC, and 49% of FoodMine). There are two main reasons behind it. First, limitations to the NLP LitSense algorithms used by FoodAtlas may limit synonyms and exhaustive tagging of the various entities, co-occurrence of entities in windows that are further away in the text body, and information that is in tables, figures, or supplementary files^59^. Second, the lack of references that are indexed and unique IDs for either foods or chemicals may introduce false positives. Further experimental validation of findings, such as the 11 novel associations with indirect evidence proposed by our link prediction pipeline, will help in accelerating the discovery and achieving completeness of the domain knowledge.

Large language models worked well in the entailment model but not for link prediction. We tested state-of-the-art language models like KG-BERT^60^ and KGLM^61^ that have better MR metrics compared to the graph-embedding models and are generalizable to unseen entities or relations^62^. For example, we obtained an MR of 191 on the validation set by fine-tuning the KG-BERT architecture with the BioBERT as a pre-trained backbone instead of the BERT, which is a significant improvement over the RotatE MR of 1,139. However, those models were not used as other metrics were significantly worse than simpler algorithms like RotatE (MRR: 0.12, hits@1: 0.08, hits@3: 0.11, and hits@10: 0.18), and training/inference time was much longer, making it infeasible to perform proper hyperparameter optimization over our large-scale FAKG. In addition, while the GPT-3.5 performance was impressive even without refinement on domain-specific data, it was not on par with the FoodAtlas pipeline, and the lack of source reference IDs defeats the purpose of one of the main pillars behind FoodAtlas: providing high-quality, trustworthy information with evidence provenance.

We identified a conflicting food-chemical relationship from the link prediction generated hypotheses. In some cases, this supports FoodAtlas’s potential to challenge established knowledge and emphasizes the necessity of experimental checking of the solid, established data. For instance, the established absence of beta-carotene synthesis in Atlantic Cod (FDC food 171955) contrasts with a high probability score (0.84 ± 0.09; **Supplementary Data 5**) of the hypothesis (Atlantic Cod, contains, beta-carotene). We sought to reconcile this through literature validation, noting that while unused genes often degrade over time due to natural selection^63^, the Atlantic Cod retains the beta,beta-carotene 15,15’-dioxygenase-like enzyme gene. Nevertheless, our further investigation considered the Atlantic Cod’s diet, particularly during its larval stage, which predominantly consists of crustaceans^64^ rich in beta-carotene^65^. Thus, there exists a plausible dietary source for beta-carotene incorporation. Additionally, we discovered literature referencing the detection of beta-carotene in commercially processed cod liver oil, albeit the exact species of cod (Gadus morhua-Atlantic Cod or others like Gadus macrocephalus-Pacific Cod) was not specified^66^.

The next version of FoodAtlas will address current limitations in data type, structure, and information source. First, we will work towards extending information extraction to the chemical concentration value in their source food (*e.g.*, cocoa contains 564 mg/serving of epicatechin)^67^. Second, not all information sources are equal, and we plan to introduce a quality score using the source trustworthiness^68^. Third, it was required that the food and chemical entities in the KG have a unique NCBI ID and PubChem ID, respectively, to ensure compatibility with existing data. Although food ontologies like FoodOn^69^ exist, we need to create and adopt a unified vocabulary of all foods and food parts that can be further extended to include processes for processed foods. Fourth, expanding FoodAtlas to capture health conditions through a UMLS ontology^70^ and dosage effects can link foods, ingredients, and their health effects in a way that can be useful for the discovery of new food-related bioactive compounds and sources^71^, food formulation and substitutions^72^, personalized diet recommendations^73^, among others. Compound information sources can also be expanded so that we include in more detail classes of molecules, such as terpenes, polyphenols, and peptides, that are of high interest^74^. Fifth, the identification and augmentation of data that the entailment model has difficulty handling, such as domain-specific hypotheses and complex sentence structures, could lead to improved performance. Finally, we will investigate further the use of pre-trained and fine-tuned large language models with active learning strategies, including those based on the network analysis indicators like centrality and modularity^75,76^, as the field has in the past few months produced striking results when it comes to efficiency, robustness, and scalability. We believe that the application of cutting-edge AI tools domains where computational science penetration has been traditionally limited, has the potential to revolutionize and pave the way for a paradigm-shift in those industries, with far reaching implications for our society and planet.

## Data Availability

All data is available at https://github.com/IBPA/FoodAtlas and http://foodatlas.ai.

## Code Availability

All code and instructions on how to reproduce the results can be found at https://github.com/IBPA/FoodAtlas.

## Acknowledgments

We would like to thank the members of the Tagkopoulos lab and Danielle Lemay from the U.S. Department of Agriculture Agricultural Research Service (USDA ARS) for helpful discussions and comments. We would also like to thank Alexis Allot from National Center for Biotechnology Information (NCBI) for running LitSense queries internally, Kyle McKillop and Kai Blumberg from the USDA ARS for providing FoodData Central (FDC) data, Anders Poulsen from the Technical University of Denmark (DTU) for providing the Frida data, Navneet Rai and Adil Muhammad from the Tagkopoulos lab for PH pair annotation, and Arielle Yoo for link prediction validation and analysis. This work was supported by the USDA-NIFA AI Institute for Next Generation Food Systems (AIFS), USDA-NIFA award number 2020-67021-32855.

## Author Contributions

J.Y. and F.L. performed all computational analyses and created figures. J.Y., F.L., and

S.K. performed the literature validation. G.S. performed the preliminary analysis. J.Y., F.L., and I.T. contributed to the critical analysis and wrote the paper. I.T. conceived and supervised all aspects of the project.

## References

1. Barabási, A.-L., Menichetti, G. & Loscalzo, J. The unmapped chemical complexity of our diet. Nat. Food 1, 33–37 (2020).

2. Elmadfa, I. & Meyer, A. L. Importance of food composition data to nutrition and public health. Eur. J. Clin. Nutr. 64, S4–S7 (2010).

3. Diana, M., Quílez, J. & Rafecas, M. Gamma-aminobutyric acid as a bioactive compound in foods: a review. J. Funct. Foods 10, 407–420 (2014).

4. Reboredo-Rodríguez, P. et al. State of the Art on Functional Virgin Olive Oils Enriched with Bioactive Compounds and Their Properties. Int. J. Mol. Sci. 18, 668 (2017).

5. Gan, J., Siegel, J. B. & German, J. B. Molecular annotation of food – Towards personalized diet and precision health. Trends Food Sci. Technol. 91, 675–680 (2019).

6. Eetemadi, A. et al. The Computational Diet: A Review of Computational Methods Across Diet, Microbiome, and Health. Front. Microbiol. 11, (2020).

7. Eetemadi, A. & Tagkopoulos, I. Methane and fatty acid metabolism pathways are predictive of Low-FODMAP diet efficacy for patients with irritable bowel syndrome. Clin. Nutr. Edinb. Scotl. 40, 4414–4421 (2021).

8. McKillop, K., Harnly, J., Pehrsson, P., Fukagawa, N. & Finley, J. FoodData Central, USDA’s Updated Approach to Food Composition Data Systems. Curr. Dev. Nutr. 5, 596 (2021).

9. Anses. Ciqual French food composition table. (2020).

10. Kapsokefalou, M. et al. Food Composition at Present: New Challenges. Nutrients 11, 1714 (2019).

11. Scalbert, A. et al. The food metabolome: a window over dietary exposure. Am. J. Clin. Nutr. 99, 1286–1308 (2014).

12. Wishart, D. FooDB Version 1.0.

13. Rakhi, N. K., Tuwani, R., Mukherjee, J. & Bagler, G. Data-driven analysis of bio-medical literature suggests broad-spectrum benefits of culinary herbs and spices. PLOS ONE 13, e0198030 (2018).

14. Afendi, F. M. et al. KNApSAcK Family Databases: Integrated Metabolite–Plant Species Databases for Multifaceted Plant Research. Plant Cell Physiol. 53, e1 (2012).

15. Duke, J. & Bogenschutz, M. J. Dr. Duke’s Phytochemical and Ethnobotanical Databases. (USDA, Agricultural Research Service Washington, DC, 1994).

16. Neveu, V. et al. Phenol-Explorer: an online comprehensive database on polyphenol contents in foods. Database 2010, bap024 (2010).

17. Rothwell, J. A. et al. Phenol-Explorer 2.0: a major update of the Phenol-Explorer database integrating data on polyphenol metabolism and pharmacokinetics in humans and experimental animals. Database 2012, bas031 (2012).

18. Rothwell, J. A. et al. Phenol-Explorer 3.0: a major update of the Phenol-Explorer database to incorporate data on the effects of food processing on polyphenol content. Database 2013, bat070 (2013).

19. Silva, A. B. da, et al. PhytoHub V1.4: A new release for the online database dedicated to food phytochemicals and their human metabolites. in np (2016).

20. White, J. PubMed 2.0. Med. Ref. Serv. Q. 39, 382–387 (2020).

21. Roberts, R. J. PubMed Central: The GenBank of the published literature. Proc. Natl. Acad. Sci. 98, 381–382 (2001).

22. Chen, X., Jia, S. & Xiang, Y. A review: Knowledge reasoning over knowledge graph. Expert Syst. Appl. 141, 112948 (2020).

23. Xu, J. et al. Building a PubMed knowledge graph. Sci. Data 7, 205 (2020).

24. Cenikj, G. et al. From language models to large-scale food and biomedical knowledge graphs. Sci. Rep. 13, 7815 (2023).

25. Harnoune, A. et al. BERT based clinical knowledge extraction for biomedical knowledge graph construction and analysis. Comput. Methods Programs Biomed. Up-date 1, 100042 (2021).

26. Diaz Gonzalez, A. D., Hughes, K. S., Yue, S. & Hayes, S. T. Applying BioBERT to Extract Germline Gene-Disease Associations for Building a Knowledge Graph from the Biomedical Literature. in 2023 the 7th International Conference on Information System and Data Mining (ICISDM) 37–42 (ACM, Atlanta USA, 2023). doi:10.1145/3603765.3603771.

27. Dang, L. D., Phan, U. T. P. & Nguyen, N. T. H. GENA: A knowledge graph for nutrition and mental health. J. Biomed. Inform. 145, 104460 (2023).

28. Haussmann, S. et al. FoodKG: A Semantics-Driven Knowledge Graph for Food Recommendation. in The Semantic Web – ISWC 2019 (eds. Ghidini, C. et al.) vol. 11779 146–162 (Springer International Publishing, Cham, 2019).

29. Ahmad, Z. et al. Active Learning Based Relation Classification for Knowledge Graph Construction from Conversation Data. in Neural Information Processing (eds. Yang, H. et al.) 617–625 (Springer International Publishing, Cham, 2020). doi:10.1007/978-3-030-63820-7_70.

30. Sun, L. et al. ASRC:A Knowledge Graph Relation Construction Model based on Active Learning and Semantic Recognition. in 2022 IEEE International Conference on Big Data (Big Data) 6025–6029 (2022). doi:10.1109/BigData55660.2022.10020502.

31. Ren, P. et al. MKGB: A Medical Knowledge Graph Construction Framework Based on Data Lake and Active Learning. in Health Information Science (eds. Siuly, S., Wang, H., Chen, L., Guo, Y. & Xing, C.) vol. 13079 245–253 (Springer International Publishing, Cham, 2021).

32. Schoch, C. L. et al. NCBI Taxonomy: a comprehensive update on curation, resources and tools. Database J. Biol. Databases Curation 2020, baaa062 (2020).

33. Allot, A. et al. LitSense: making sense of biomedical literature at sentence level. Nucleic Acids Res. 47, W594–W599 (2019).

34. Lee, J. et al. BioBERT: a pre-trained biomedical language representation model for biomedical text mining. Bioinformatics btz682 (2019) doi:10.1093/bioinformatics/btz682.

35. Devlin, J., Chang, M.-W., Lee, K. & Toutanova, K. BERT: Pre-training of Deep Bidirectional Transformers for Language Understanding. Preprint at 10.48550/arXiv.1810.04805 (2019).

36. Liu, Y., et al. RoBERTa: A Robustly Optimized BERT Pretraining Approach. Pre-print at 10.48550/arXiv.1907.11692 (2019).

37. Brown, T. et al. Language Models are Few-Shot Learners. Adv. Neural Inf. Process. Syst. 33, 1877–1901 (2020).

38. National Food Institute, Technical University of Denmark. Food data (frida.fooddata.dk), version 4.2, 2022.

39. Rossi, A., Barbosa, D., Firmani, D., Matinata, A. & Merialdo, P. Knowledge Graph Embedding for Link Prediction: A Comparative Analysis. ACM Trans. Knowl. Discov. Data 15, 14:1–14:49 (2021).

40. Ali, M. et al. PyKEEN 1.0: a Python library for training and evaluating knowledge graph embeddings. J. Mach. Learn. Res. 22, 82:3723–82:3728 (2021).

41. Bordes, A., Usunier, N., Garcia-Duran, A., Weston, J. & Yakhnenko, O. Translating Embeddings for Modeling Multi-relational Data. in Advances in Neural Information Processing Systems vol. 26 (Curran Associates, Inc., 2013).

42. Dong, X. et al. Knowledge vault: a web-scale approach to probabilistic knowledge fusion. in Proceedings of the 20th ACM SIGKDD international conference on Knowledge discovery and data mining 601–610 (Association for Computing Machinery, New York, NY, USA, 2014). doi:10.1145/2623330.2623623.

43. Yang, B., Yih, W., He, X., Gao, J. & Deng, L. Embedding Entities and Relations for Learning and Inference in Knowledge Bases. Preprint at 10.48550/arXiv.1412.6575 (2015).

44. Ji, G., He, S., Xu, L., Liu, K. & Zhao, J. Knowledge Graph Embedding via Dynamic Mapping Matrix. in Proceedings of the 53rd Annual Meeting of the Association for Computational Linguistics and the 7th International Joint Conference on Natural Language Processing (Volume 1: Long Papers) 687–696 (Association for Computational Linguistics, Beijing, China, 2015). doi:10.3115/v1/P15-1067.

45. Trouillon, T., Welbl, J., Riedel, S., Gaussier, E. & Bouchard, G. Complex Embeddings for Simple Link Prediction. in Proceedings of The 33rd International Conference on Machine Learning 2071–2080 (PMLR, 2016).

46. Sun, Z., Deng, Z.-H., Nie, J.-Y. & Tang, J. RotatE: Knowledge Graph Embedding by Relational Rotation in Complex Space. Preprint at http://arxiv.org/abs/1902.10197 (2019).

47. Youn, J. & Tagkopoulos, I. KGLM: Integrating Knowledge Graph Structure in Language Models for Link Prediction. Preprint at 10.48550/arXiv.2211.02744 (2022).

48. Kim, S. et al. PubChem Protein, Gene, Pathway, and Taxonomy Data Collections: Bridging Biology and Chemistry through Target-Centric Views of PubChem Data. J. Mol. Biol. 434, 167514 (2022).

49. Hooton, F., Menichetti, G. & Barabási, A.-L. Exploring food contents in scientific literature with FoodMine. Sci. Rep. 10, 16191 (2020).

50. Reifen, R., Karlinsky, A., Stark, A. H., Berkovich, Z. & Nyska, A. α-Linolenic acid (ALA) is an anti-inflammatory agent in inflammatory bowel disease. J. Nutr. Biochem. 26, 1632–1640 (2015).

51. Singer, P. et al. Effects of dietary oleic, linoleic and alpha-linolenic acids on blood pressure, serum lipids, lipoproteins and the formation of eicosanoid precursors in patients with mild essential hypertension. J. Hum. Hypertens. 4, 227–233 (1990).

52. Lawrence, G. D. & Sawyer, D. T. The chemistry of biological manganese. Coord. Chem. Rev. 27, 173–193 (1978).

53. Schramm, V. L. Manganese in Metabolism and Enzyme Function. (Elsevier, 2012).

54. Aguirre, J. D. & Culotta, V. C. Battles with Iron: Manganese in Oxidative Stress Protection *. J. Biol. Chem. 287, 13541–13548 (2012).

55. Nagao, A., Maeda, M., Lim, B. P., Kobayashi, H. & Terao, J. Inhibition of β-carotene-15,15′-dioxygenase activity by dietary flavonoids. J. Nutr. Biochem. 11, 348–355 (2000).

56. Garcia-Mas, J. et al. The genome of melon ( *Cucumis melo* L.). Proc. Natl. Acad. Sci. 109, 11872–11877 (2012).

57. Joradon, P. et al. Ergosterol Content and Antioxidant Activity of Lion’s Mane Mushroom (Hericium erinaceus) and Its Induction to Vitamin D2 by UVC-Irradiation: in *Proceedings of the 8th International Conference on Agricultural and Biological Sciences* 19–28 (SCITEPRESS - Science and Technology Publications, Shenzhen, China, 2022). doi:10.5220/0011594600003430.

58. Sun, Y., Nzekoue, F. K., Vittori, S., Sagratini, G. & Caprioli, G. Conversion of ergosterol into vitamin D2 and other photoisomers in Agaricus bisporus mushrooms under UV-C irradiation. Food Biosci. 50, 102143 (2022).

59. Herzig, J., Nowak, P. K., Müller, T., Piccinno, F. & Eisenschlos, J. M. TAPAS: Weakly Supervised Table Parsing via Pre-training. in Proceedings of the 58th Annual Meeting of the Association for Computational Linguistics 4320–4333 (2020). doi:10.18653/v1/2020.acl-main.398.

60. Yao, L., Mao, C. & Luo, Y. KG-BERT: BERT for Knowledge Graph Completion. Preprint at 10.48550/arXiv.1909.03193 (2019).

61. Youn, J. & Tagkopoulos, I. KGLM: Integrating Knowledge Graph Structure in Language Models for Link Prediction. in Proceedings of the 12th Joint Conference on Lexical and Computational Semantics (*SEM 2023) (eds. Palmer, A. & Camacho-collados, J.) 217–224 (Association for Computational Linguistics, Toronto, Canada, 2023). doi:10.18653/v1/2023.starsem-1.20.

62. Zha, H., Chen, Z. & Yan, X. Inductive Relation Prediction by BERT. Proc. AAAI Conf. Artif. Intell. 36, 5923–5931 (2022).

63. Albalat, R. & Cañestro, C. Evolution by gene loss. Nat. Rev. Genet. 17, 379–391 (2016).

64. Hamre, K. Nutrition in cod (Gadus morhua) larvae and juveniles. ICES J. Mar. Sci. 63, 267–274 (2006).

65. Maoka, T. Carotenoids in Marine Animals. Mar. Drugs 9, 278–293 (2011).

66. Luterotti, S., Franko, M. & Bicanic, D. Ultrasensitive determination of β-carotene in fish oil-based supplementary drugs by HPLC-TLS. J. Pharm. Biomed. Anal. 21, 901– 909 (1999).

67. Crozier, A., Jaganath, I. B. & Clifford, M. N. Dietary phenolics: chemistry, bioavailability and effects on health. Nat. Prod. Rep. 26, 1001–1043 (2009).

68. Kyngäs, H., Kääriäinen, M. & Elo, S. The Trustworthiness of Content Analysis. in The Application of Content Analysis in Nursing Science Research (eds. Kyngäs, H., Mikkonen, K. & Kääriäinen, M.) 41–48 (Springer International Publishing, Cham, 2020). doi:10.1007/978-3-030-30199-6_5.

69. Dooley, D. M. et al. FoodOn: a harmonized food ontology to increase global food traceability, quality control and data integration. Npj Sci. Food 2, 23 (2018).

70. Bodenreider, O. The Unified Medical Language System (UMLS): integrating biomedical terminology. Nucleic Acids Res. 32, D267–D270 (2004).

71. Min, W., Liu, C., Xu, L. & Jiang, S. Applications of knowledge graphs for food science and industry. Patterns 3, 100484 (2022).

72. Ławrynowicz, A., Wróblewska, A., Adrian, W. T., Kulczyński, B. & Gramza-Michałowska, A. Food Recipe Ingredient Substitution Ontology Design Pattern. Sensors 22, 1095 (2022).

73. Chen, Y., Subburathinam, A., Chen, C.-H. & Zaki, M. J. Personalized Food Recommendation as Constrained Question Answering over a Large-scale Food Knowledge Graph. in Proceedings of the 14th ACM International Conference on Web Search and Data Mining 544–552 (ACM, Virtual Event Israel, 2021). doi:10.1145/3437963.3441816.

74. Degtyarenko, K. et al. ChEBI: a database and ontology for chemical entities of biological interest. Nucleic Acids Res. 36, D344–D350 (2008).

75. Rodrigues, F. A. Network Centrality: An Introduction. in A Mathematical Modeling Approach from Nonlinear Dynamics to Complex Systems (ed. Macau, E. E. N.) 177– 196 (Springer International Publishing, Cham, 2019). doi:10.1007/978-3-319-78512-7_10.

76. Wagner, G. P., Pavlicev, M. & Cheverud, J. M. The road to modularity. Nat. Rev. Genet. 8, 921–931 (2007).

77. Abdel-Salam, O. M. E. et al. Cannabis-induced impairment of learning and memory: effect of different nootropic drugs. EXCLI J. 12, 193–214 (2013).

78. Steigerwalt, R. D. et al. Mirtogenol potentiates latanoprost in lowering intraocular pressure and improves ocular blood flow in asymptomatic subjects. Clin. Ophthalmol. Auckl. NZ 4, 471–476 (2010).

79. Zhang, G. et al. Therapeutic Efficiency of an External Chinese Herbal Formula of Mammary Precancerous Lesions by BATMAN-TCM Online Bioinformatics Analysis Tool and Experimental Validation. Evid.-Based Complement. Altern. Med. ECAM 2019, 2795010 (2019).

80. Bernier, C., Goetz, C., Jubinville, E. & Jean, J. The New Face of Berries: A Review of Their Antiviral Proprieties. Foods 11, 102 (2021).

81. Monroig, Ó. et al. Biosynthesis of Polyunsaturated Fatty Acids in Octopus vulgaris: Molecular Cloning and Functional Characterisation of a Stearoyl-CoA Desaturase and an Elongation of Very Long-Chain Fatty Acid 4 Protein. Mar. Drugs 15, 82 (2017).

82. Schultz, I. R., Skillman, A. & Woodruff, D. Domoic acid excretion in dungeness crabs, razor clams and mussels. Mar. Environ. Res. 66, 21–23 (2008).

83. Yazdanparast, R. & Alavi, M. Antihyperlipidaemic and antihypercholesterolaemic effects of Anethum graveolens leaves after the removal of furocoumarins. Cytobios 105, 185–191 (2001).

84. Kruse, A. B. et al. What is the impact of the adjunctive use of omega-3 fatty acids in the treatment of periodontitis? A systematic review and meta-analysis. Lipids Health Dis. 19, 100 (2020).

