## Supplementary Information for "FoodAtlas: Automated Knowledge Extraction of Food and Chemicals from Literature"

### Table of Contents

1 Supplementary Text ... 4

1.1 Additional analysis ... 4

1.1.1 Percentage of associations with literature citations in FooDB ... 4

1.1.2 Estimation of the total number of triplets in PubMed Central ... 4

1.1.3 Lit2KG generated triplets vs. external database triplets ... 5

1.1.4 Validation using FoodMine ... 5

1.1.5 Active learning performance ... 7

1.1.6 Understanding the entailment predictions ... 8

1.1.7 Entailment model performance comparison across literature sections ... 8

1.1.8 Validation for link prediction ... 9

1.1.9 Benchmark with FooDB. ... 15

1.2 Methods ... 16

1.2.1 Food name collection and LitSense query ... 16

1.2.2 Data annotation ... 17

1.2.3 Entailment model ... 18

1.2.4 Link prediction ... 22

1.3 FoodAtlas Knowledge Graph ... 24

1.3.1 Entity types ... 24

### 1 Supplementary Text

#### 1.1 Additional analysis

##### 1.1.1 Percentage of associations with literature citations in FooDB

We downloaded *foodb_2020_04_07_csv/Content.csv* from the FooDB (*https://foodb.ca*) website and analyzed *citation_type* and *citation* columns, which represent a generalized description and a more specific identifier for a citation of a given food-chemical association. Among 5,145,532 FooDB associations, 3,273,562 are classified as *PREDICTED* by *PATHBANK* or *HMDB*. 955,962 associations are noted as *MANUAL* without a citation. Finally, 895,467 associations are linked to other databases without specific references to scientific literature. Therefore, approximately 20,541 associations are with a scientific literature reference in FooDB, which is 0.4% of the total number of associations.

##### 1.1.2 Estimation of the total number of triplets in PubMed Central

To estimate all food-chemical associations available to extract in scientific literature, we extrapolated the total number of triplets extractable from PubMed using the statistics of the FoodAtlas knowledge graph (FAKG). First, we estimated the expected number of unique triplets extracted from each publication, denoted as *m*, by having the unique number of triplets extracted from PH pairs in FAKG divided by the unique number of PMIDs. While calculating, since most triplets resulted from FoodAtlas entailment predictions, we weighted each triplet based on the prediction probabilities,

$$m = \frac{1}{|L|}\sum_{t \in T} 1 \cdot p_t = \frac{1}{146{,}507} \cdot 193{,}310.42 = 1.32,$$

where $L$ is the set of PMIDs in FAKG, $T$ is the set of triplets, and $p_t$ is the entailment probability for the triplet $t$. Then, we counted the number of papers relevant to food in PubMed using the advanced search query "(food) OR (fruit) OR (vegetable)", which returned 1,588,596 unique PMIDs. With this information, we estimated that our pipeline would be able to extract 1.32 x 1,588,596 ≈ 2.1 M triplets from unstructured text alone, not including tables, supplementary files, etc.

###### 1.1.3 Lit2KG generated triplets vs. external database triplets

The Lit2KG pipeline integrated, on average, 39% of the contains triplets from the external databases (323 out of 529 FDC triplets, 371 out of 1,055 Phenol-Explorer triplets, and 1,873 out of 8,375 Frida triplets were also discovered by Lit2KG pipeline) (**Fig. 2d**). We also identified 219,854 medium-quality triplets that were not reported in external databases (**Fig. 2e**).

###### 1.1.4 Validation using FoodMine

To evaluate the chemical coverage of FAKG, we compared chemicals in FAKG to that in FoodMine[1], a database exhaustively curating chemicals for cocoa and garlic from the literature. In this section, we describe the methodology of the validation.

###### 1.1.4.1 Processing chemicals in FoodMine

FoodMine initially contained 598 and 289 unique chemicals for cocoa and garlic, respectively. However, due to different chemical indexing methods, the chemicals in FoodMine were not directly comparable to those in FAKG. In the FoodMine paper, the authors describe their method for chemical disambiguation using *structural code*, *i.e.*, the first 14 characters of InChIKey[1]. To be comparable, we follow the same approach as

FoodMine did. Since every chemical in FAKG has a PubChem CID, we retrieved InChiKey for each chemical and derived its structural code. Finally, we dropped chemicals without structural code in FoodMine, resulting in 301 and 176 unique chemicals for cocoa and garlic, respectively.

###### 1.1.4.2 Processing chemicals in FAKG

The processing required for FAKG was more complicated than that for FoodMine. There were four types of evidence in FAKG: entailment annotation, entailment prediction, link prediction, and external databases. We treated entailment annotation and external database evidence as positive evidence supporting a food containing a chemical. For entailment prediction, if we filtered by the NCBI Taxonomy ID, FAKG had 1,289 and 1,376 chemicals with unique PubChem CIDs for cocoa and garlic, respectively. Then, as mentioned in the section above, we retrieved the structural code for each chemical, resulting in 499 and 691 unique structural codes for cocoa and garlic, respectively. For a fair comparison between FAKG and FoodMine, we manually annotated and excluded the false positives and NER errors of entailment prediction. To do that, we collected all the entailment prediction evidence in FAKG for all structural codes, resulting in 5,506 and 9,616 pieces of evidence for cocoa and garlic, respectively (**Supplementary Data 6**). Note that LitSense also tagged “chocolate” as cocoa, so 5,506 evidence were dropped. In this case, we were only interested in the existence of entailment prediction evidence regarding cocoa or garlic containing a structural code. Thus, once we encountered one piece of positive evidence, we skipped the rest of the evidence associated with that structural code, which means we discarded a structural code only if all evidence was false positive. Eventually, FAKG contained 366 and 400 unique structure codes for cocoa and garlic, respectively, supported by the entailment prediction evidence. For link prediction evidence, we annotated the structural codes following **Section 1.1.8 of Supplementary Information**, which resulted in 14 and 8 unique chemicals for cocoa and garlic, respectively (**Supplementary Data 7**). After removing the duplicated structural codes between entailment and link prediction, we had 379 and 406 chemicals with unique structural codes.

###### 1.1.4.3 Limitations

There were limitations in this comparison. First, FoodMine only curates quantified chemicals, while FAKG curates identified food chemicals, regardless of the existence of quantification. Second, we adopted the same methodology of FoodMine to match the chemicals in FoodMine and FAKG. However, the method considered chemicals within the same isomeric species identical by only looking at the segment InChIKeys. Lastly, as mentioned in the last section, FAKG focuses on raw food, while FoodMine does not discriminate between raw and non-raw food.

##### 1.1.5 Active learning performance

We calculated the performance of the entailment model at the final AL round (r = 10) by averaging all 400 models (100 random seeds for each of the 4 AL strategies trained on the same data). Compared to the initial round (r = 1), precision at the final round (r = 10) increased by 10.8% (0.74 ± 0.04 vs. 0.82 ± 0.03, respectively, p-value = 1.3 × 10-155), recall by 21.7% (0.69 ± 0.11 vs. 0.84 ± 0.07, respectively, p-value = 3.7 × 10-89), and F1 score by 16.9% (0.71 ± 0.05 vs. 0.83 ± 0.03, respectively, p-value = 2.8 × 10-206) (**Supplementary Fig. 6** and **Supplementary Table 1**).

##### 1.1.6 Understanding the entailment predictions

**Table 1** shows selected entailment model predictions for each of the four outcome scenarios, true positive (TP), false positive (FP), true negative (TN), and false negative (FN), of the confusion matrix. For certain TP and TN ($p \approx 1$ and $p \approx 0$, respectively), PH pairs are usually straightforward, such as premises with simple sentence structure (index 1) or hypotheses with wrong food and food part matching (index 3). For certain FP and FN and uncertain predictions, we found that premises often require the models to use domain expertise to predict correctly (e.g., *essential nutrient* implies chemicals may not naturally occur in species as in index 5) or are titles of the paper that are hypotheses yet to be validated (index 6). Uncertain predictions ($p \approx 0.5$) had a higher standard deviation of the probability scores assigned by the 400 entailment models at the final round than the certain predictions (**Table 1**).

##### 1.1.7 Entailment model performance comparison across literature sections

We compared how sentences from different literature sections affected the production entailment model performance. Since the production model no longer had a designated test set, we performed 10-fold cross-validation to get the prediction score for each PH pair using the same train and validation splits and best hyperparameter described in **Section 1.2.3.2 of Supplementary Information**. To further increase the robustness, we repeated the cross-validation 50 times with different weight initializations such that the prediction score for each PH pair was the average of 50 runs. **Supplementary Table 2** summarizes the confusion matrices for each section. We found that for the PH pairs from the introduction section, the model achieved the highest precision, F1, and the third-best recall scores. Also, regarding prediction scores, the model predicted the PH pairs of the introduction section significantly differently from those of the other sections (vs. Discussion, $p$-value = $7.8 \times 10^{-33}$; vs. Abstract, $p$-value = $3.2 \times 10^{-104}$; vs. Methods, $p$-value = $6.7 \times 10^{-206}$; vs. Results, $p$-value = $1.8 \times 10^{-13}$; vs. Conclusion, $p$-value = $9.4 \times 10^{-4}$; vs. Title, $p$-value = $4.3 \times 10^{-18}$; vs. Table, $p$-value = $1.1 \times 10^{-23}$; vs. Others, $p$-value = $1.0 \times 10^{-34}$. $p$-values were computed by the two-sided t-test with the Benjamini-Hochberg adjustment[2]).

##### 1.1.8 Validation for link prediction

To validate the utility of link prediction in discovering novel food-chemical relations, we manually annotated the triplets predicted by the best link prediction (LP) model. Because the inputs to LP were merely triplets (i.e., no longer using premise-hypothesis pairs), finding and annotating the evidence for link prediction triplets proved challenging without domain expertise. Therefore, we seek help from a postdoctoral biochemistry expert to annotate the triplets for LP validation. For this section, we will discuss the procedure of query, search, and annotation.

###### 1.1.8.1 Annotation datasets

To validate that the link predictor was well-calibrated, we annotated all 466 triplets with probability score greater than 90% as well as 200 randomly sampled triplets with 20 triplets from each of the 10 equally spaced bins (**Supplementary Data 5**). The reason for selecting these two sets was because the former would allow the discovery of novel food-chemical associations by looking at the most likely triplets, while the latter would show how well our model is calibrated based on a scrutinized literature search. However, note that we skipped some unwanted triplets, including triplets with entities with LitSense NER Error and triplets with chemical entities that were chemical groups rather than a specific chemical structure.

###### 1.1.8.2 Query standard

We searched four sources, which are PubChem taxonomy[3], Bing Chat, Google Scholar, and Google. We queried PubChem taxonomy and Bing Chat because they returned more straightforward responses compared to Google Scholar and Google, which had better coverage.

For PubChem taxonomy, we queried the scientific name of the food entity of the triplet to retrieve the evidence. Since PubChem taxonomy retrieved evidence based on fuzzy search[3], we reviewed and verified the evidence.

For Bing Chat, we queried the search engine by prompting, “Does *Theobroma cacao* contain stearic acid?” where synonyms for foods and chemicals were also used to query independently. Specifically, for food entities, we used all the synonyms listed in the NCBI taxonomy, while for chemical entities, we tried a subset of synonyms that were most commonly used (**Supplementary Fig. 11**). We checked the evidence links returned by Bing Chat and verified their correctness.

For Google Scholar and Google, we formulated the query by concatenating the food and chemical names, e.g., *Theobroma cacao* stearic acid. The same treatment for synonyms performed in Bing Chat was applied.

###### 1.1.8.3 Annotation standard

If we found the evidence, we annotated the triplets as *Yes*, and the reference sources were recorded (**Supplementary Data 5**). The triplets were annotated as *Unknown* if we found no evidence since we could not conclude that the corresponding triplet was novel without exhaustively searching the entire internet. Finally, a few triplets were annotated as *No,* if food could not biologically contain a chemical. For example, a triplet with the chemical *1,1,1-Trichloroethane* was labeled as *No* because the chemical could only be synthesized in labs.

###### 1.1.8.4 Validation for *Unknown* triplets

*Unknown* triplets, described in the last section, not being found on the internet could be either novel or wrong. Since experimentally validating triplets would be expensive, we describe a heuristic to decide if a triplet is likely true based on its metabolic and evolutionary viability, where we consider a triplet is likely to be true if it passes at least one of the tests. We start this section with a description of our tests, followed by a specific discussion for each triplet that has passed the test.

**Metabolic viability test**. A chemical is likely to present in a species if a metabolic pathway exists within the species to produce that chemical, and we utilized the following approaches to identify the pathways likely to exist within species. First, we checked if a species can synthesize enzymes known to produce the chemical of interest. To do this, we searched for the enzymes of the chemical in databases, like KEGG[4], and literature. Then, we retrieved the enzyme amino acid sequence from UniProt[5]. Finally, we input the retrieved enzyme sequences and the species of interest in the NCBI BLAST[6] query to find the homologous enzymes with similar sequences producible by the species. Based on the assumption that similar sequences share a similar function, we considered the triplet passing the metabolomic viability test if the searched enzymes were homologous with the query sequence by hitting the *E*-value of 0, implying that the observed similarity between the query and subject sequences was highly unlikely to have arisen randomly or by chance. Note that not all chemical synthesis requires enzymes. In those cases, we looked for the chemical equation that produced the chemical and verified whether the required reactants and catalysts were found in the species. If found, we also considered the triplet passing the test.

**Evolutionary viability test**. The metabolic viability test could be challenging for some species because they were not found in KEGG or BLAST. In such cases, we resorted to an evolutionary viability test. Specifically, a triplet is considered to pass the test if there are any species under the same genus of the species of interest containing the chemical. For example, we could not find any metabolic pathway for phenol in *Cinnamomum aromaticum* (Chinese cinnamon), but we found phenol in *Cinnamomum zeylanicum* (Ceylon cinnamon)[7], and hence the triplet passed the evolutionary viability test.

**Metabolically viable triplet 1: *Hericium erinacerus* (bearded tooth) contains lumisterol**. We found a recent source indicating that lumisterol can be naturally synthesized with ergosterol by UV irradiation (e.g., sunlight)[8], where the bearded tooth is known to contain ergosterol[9].

**Metabolically viable triplet 2: *Cicer arietinum* (chickpea) contains triglyceride**. Through a literature search, we found an enzyme, phospholipid:diacylglycerol acyltransferase, that could synthesize triacylglycerol (synonym for triglyceride)[10], and found its protein sequence (UniProt ID: Q9FNA9) for *Arabidopsis thaliana* (mouse-ear cress). Through BLAST (blastp), four phospholipid:diacylglycerol acyltransferase-like enzymes in chickpeas with zero *E*-value and 78.4%, 76.1%, 76.7%, and 56.9% percentage of identity (RefSeq ID: XP_004510434.1, XP_004507978.1, XP_012573333.1, and XP_004506121.1) were retrieved.

**Metabolically viable triplet 3: *Cuminum cyminum* (cumin) contains sodium caffeate**. Sodium caffeate can be synthesized naturally by caffeic acid and sodium through a neutralization process without enzymes. Caffeic acid can be found in cumin[11] while sodium is present in all plants.

**Metabolically viable triplet 4: *Gadus morhua* (Atlantic cod) contains beta-carotene**. Through a literature search, we found beta-carotene-15,15'-dioxygenase, an enzyme in *Homo sapiens* (human) that catalyzes the reaction involved in beta-carotene[12]. We used the corresponding protein sequence (UniProt ID: Q9HAY6) and identified two protein sequences through BLAST (blastp) with zero E-values and percentage of identities of 58.5% and 53.9% respectively (RefSeq ID: XP_030205463.1 and XP_030208886.1).

**Evolutionarily viable triplet 1: *Cucumis melo var. dudaim* (Dudaim melon) contains matairesinol**. Secoisolariciresinol dehydrogenase is an enzyme that can produce matairesinol and is found in various plants[13]. Although we could not find a source that directly indicates *Cucumis melo var. dudaim* contains this enzyme, we found that *Cucumis melo* (muskmelon) and *Cucumis melo var. makuwa* (oriental melon), two species sharing the same genus with *Cucumis melo var. dudaim*, contain secoisolariciresinol dehydrogenase (UniProt ID: A0A1S3CT49 and A0A5A7UNV0). Since the species *Cucumis melo* and a variant *Cucumis melo var. makuwa* (oriental melon) that shares the same parent subspecies (*Cucumis melo subsp. agrestis*) as *Cucumis melo var. dudaim* contains secoisolariciresinol dehydrogenase, likely, *Cucumis melo var. dudaim* and all *Cucumis melo*'s subspecies and variants contain this enzyme.

While there is a reference genome in BLAST for *Cucumis melo var. dudaim*, currently the only source for this sequencing data analyzes whole chloroplast gene assembly and is currently unpublished (https://www.ncbi.nlm.nih.gov/nuccore/1240947550), meaning that the proteome and genome are currently incomplete. Additionally, using BLAST (TBLASTN) to search for the DNA sequence for secoisolariciresinol dehydrogenase (RefSeq ID: XP_008467261.1) in *Cucumis melo*, we found the top two non-predicted sequences with E-values of $10^{-180}$ and percent identities of 96.48% (RefSeq ID: LN681895.1 and LN713263.1). The first reference sequence is a scaffold, but the second says it is located on chromosome 9 of the *Cucumis melo* nuclear genome, and the chloroplast genome has little similarity to the nuclear genome, especially chromosome 9[14]. Therefore, given the current sequencing data for *Cucumis melo var. dudaim*, we are not able to find secoisolariciresinol dehydrogenase because the current sequencing data is incomplete and the current genomic data does not contain the region that has secoisolariciresinol dehydrogenase.

**Evolutionarily viable triplet 2: *Cinnamomum aromaticum* (Chinese cinnamon) contains phenol**. Phenol can be found in *Cinnamomum zeylannicum* (Ceylon cinnamon)[7], which shares the same genus, *Cinnamomum*, with Chinese cinnamon.

**5 non-passing triplets**. Diospyros kaki (Japanese persimmon) contains 3-Rhamnosyl-Glucosyl Quercetin, Anthriscus cerefolium (chervil) contains loxoprofen, Allium schoenoprasum (chive) contains salicylic acid, Juglans cinerea (butternut) contains lawsone, and Curcuma longa (turmeric) contains 3-Rhamnosyl-Glucosyl Quercetin.

###### 1.1.9 Benchmark with FooDB.

We benchmarked with FooDB, the existing state-of-the-art food resource which aggregates many other popular food databases, and by comparing with FooDB, we would be able estimate the overall coverage of FoodAtlas with regard to all the existing food sources. **Supplementary Fig. 5** shows the comparison between FoodAtlas and FooDB. For the two databases to be comparable, we only considered foods and chemicals associated with NCBI Taxonomy ID (NCBI ID) and PubChem CID.

For FoodAtlas, we first extracted all the triplets with *contains* relation, resulting in 243,231 unique triplets. Within them, we counted 536 and 11,908 unique NCBI IDs and CIDs, forming 126,082 triplets with unique NCBI ID-CID pairs. The count of unique pairs was smaller than the number of triplets because different food parts were considered different triplets while sharing the same NCBI ID. Among them, 106,082 are exclusively included by FoodAtlas. To further investigate the distribution of quality of the 106,082 triplets, we retrieved all the evidence stored in FoodAtlas for the associated triplets and assigned each triplet with the highest evidence quality. For example, if a triplet had high- and medium-quality evidence simultaneously, the triplet was considered a high-quality triplet. This gave us 2,091 high-, 94,095 medium-, and 9,896 low-quality triplets, adding to 106,082.

For FooDB, we downloaded the entire database with 5,007,500 food-chemical associations. Out of 992 original foods, 600 unique NCBI IDs were found. For 70,477 original chemicals in FooDB, however, CIDs were not reported in the downloadable content. To accommodate, we used InChIKeys reported in FooDB to identify the corresponding CIDs with PucChem Identifier Exchange Service (https://pubchem.ncbi.nlm.nih.gov/docs/identifier-exchange-service). This process returned 64,125 unique CIDs in FooDB. We then extracted food-chemical pairs that were associated with using these 600 and 64,125 NCBI IDs and CIDs in FooDB, resulting in 2,480,768 associations out of the original 5,007,500. After dropping NCBI IDs and CIDs without any association, FooDB had 598 and 51,130 unique NCBI IDs and CIDs, forming 2,480,768 associations.

To make a more fine-grained comparison, we also evaluated FooDB without associations predicted via metabolic pathways. This included associations imported from PathBank[15] and HMDB[16]. This resulted in 651,680 unique associations compared to the previous 2,480,768.

#### 1.2 Methods

##### 1.2.1 Food name collection and LitSense query

We collected 650 unique NCBI Taxonomy IDs (NCBI IDs) from FooDB (*https://foodb.ca*), FDC[17], Phenol-Explorer[18–20], and Frida (*https://frida.fooddata.dk*). Note that the FoodAtlas framework requires each food item to have an NCBI ID; thus, we discarded any food items in these databases without NCBI IDs. After downloading the entire NCBI Taxonomy database from the FTP site (*https://ftp.ncbi.nlm.nih.gov/pub/taxonomy/new_taxdump; accession time: 9:53 AM on November 30th*, 2022), we found the common name, Genbank name, and scientific name entries of the 650 NCBI IDs in the downloaded file, *names.dmp*, which resulted in 1,959 food names.

###### 1.2.2 Data annotation

We deployed an annotation platform with a graphical user interface on Label Studio. We had two annotators labeled each premise-hypothesis (PH) pair as one of *entails* (i.e., the premise supports the hypothesis), *does not entail* (i.e., the premise does not support the hypothesis), and *skip* (which will be defined later in the paragraph). The annotators did a preliminary annotation session and reported some observations, which we used to create basic annotation standards for annotators to follow for the following annotation sessions. We decided that for a PH pair, (a) if LitSense returned an incorrect NCBI Taxonomy ID or MeSH ID for the hypothesis, then the PH pair should be *skip*, and (b) the annotation should not rely on sources other than premise, i.e., even though it is well-known that lemon contains vitamin C, but if the premise does not support the claim, the PH pair should be *does not entail*. Note that we desired the annotation standard to be as general as possible to avoid biasing the subsequent training of the entailment model. We ensured the annotation quality by only using the PH pairs whose annotated labels were the consensus among annotators.

##### 1.2.3 Entailment model

###### 1.2.3.1 Model configuration

We used BioBERT[21], a pre-trained language model pre-trained on the biomedical corpus, and the English Wikipedia and BooksCorpus, as our entailment model. Specifically, we imported the pre-trained model, dmis-lab/biobert-v1.1, from HuggingFace[22], with the default setup, except that we changed the pad token type ID to 1 to accommodate sentence-pair classification schema. We implemented the entailment model with PyTorch[23], a deep-learning framework. We performed the grid search hyperparameter tuning over the batch size, learning rate of the AdamW[24] optimizer, and the number of epochs. When an input sentence pair exceeded the maximum length of 512, we truncated the longer sentence, which was always the first sentence, *i.e.*, premise. We relied on four Nvidia RTX A5000 GPUs for the training and inference.

###### 1.2.3.2 Production model

Once we completed the evaluation for entailment models with different active learning strategies, we deployed the production entailment model, where we used the entire labeled dataset (*i.e.*, training, validation, and test sets) to train it. Then, to configure the optimal hyperparameters for the production model, we performed a 10-fold cross-validation, where each fold contained a unique set of premises and a unique set of hypotheses. Again, we used the same grid search space, i.e., learning rate = {$2x10^{-5}$, $5x10^{-5}$}, epochs = {3, 4}, and batch size = {16, 32}, described in the main method section, where the one with the best mean precision across ten folds was chosen for the production model. Finally, using the best hyperparameter set, we trained our production model, an ensemble system, by training 100 entailment models with different random initializations using the entire dataset (*i.e.*, all ten folds). Given one input premise-hypothesis (PH) pair, the prediction of the production model was then the mean of predicted scores of the 100 entailment models.

###### 1.2.3.3 Active learning

The available scientific literature online is too large to annotate manually. Thus, we experimented with active learning, a subfield of machine learning which aims the models to choose the training data to be labeled for the models to achieve better performance with less labeled data[25]. To compare different active learning strategies without overly burdensome manual annotation labor, we employed a *pool-based active learning* scenario, which assumes a small subset of labeled data exists in a closed pool of data, allowing the model to select new training samples from the pool[25]. Here, we describe more details regarding each active learning strategy.

**Maximum Likelihood Active Learning.** The maximum likelihood active learning continuously sampled the PH pairs with the most certainly positive probability scores. Because we only added the positives to the knowledge graph, so we hypothesized that the knowledge graph would grow the fastest with this active learning. Specifically, we sorted unsampled PH pairs based on their predicted probability scores. We then added the highest 412 PH pairs to the sampled data for the next active learning round training.

**Maximum Entropy Active Learning.** The maximum entropy active learning sampled the PH pairs that the model was most uncertain about. The PH pairs with uncertain predictions were close to the decision boundary of the model, so we hypothesized that the entailment model would benefit from learning these data. For binary classification, this means,

$$uncertainty = \min(1 - p, p),$$

where $p$ is the probability score associated with the entailment prediction. We can observe that the uncertainty score is the highest when $p = 0.5$.

**Stratified Active Learning.** The stratified active learning split the data pool into ten bins of equal interval (i.e., [0, 0.1], [0.1, 0.2), ..., [0.9, 1.0)), and the entailment models selected the same number of samples from each bin randomly. However, especially for the late active learning rounds, the number of samples in a bin might be less than the number of samples the model required to draw. Therefore, we ensured to train the model with the same amount of data in each round by using the following simple algorithm for each stratified active learning round sampling:

1. Initialize the number of bins B = 10 and the total number of samples drawn per round N = 412.
2. Split the data pool into B equal-interval bins.
3. Check if, for all $B$ bins, there exists at least (N / B) samples:
    a. If true, randomly draw (N / B) samples from each bin.
    b. If false, update B ← B - 1, and start from step 2.

**Random Sampling.** Random sampling is our baseline equivalent to no active learning. Every PH pair has an equal chance to be sampled by the model.

###### 1.2.3.4 Pseudocode

**Algorithm 1.** Training an entailment model with pool-based active learning (AL).

**Input:** $\{\boldsymbol{X_A}, \boldsymbol{y_A}, \boldsymbol{X_T}, \boldsymbol{H}\}$, where $\boldsymbol{X_A}$ is a set of annotated PH pairs for training the model, $\boldsymbol{y_A}$ is an array of labels of $\boldsymbol{X_A}$, $\boldsymbol{X_T}$ is a set of PH pairs predicted by the trained model, and $\boldsymbol{H}$ is the hyperparameter search space.
**Output:** $\{\boldsymbol{X_T}, \widehat{\boldsymbol{y_T}}\}$, where $\boldsymbol{X_T}$ is the same as the input, and $\widehat{\boldsymbol{y_T}}$ is an array of labels of $\boldsymbol{X_T}$ predicted by the model across multiple AL rounds.

$(\boldsymbol{X_{train}}, \boldsymbol{y_{train}}), (\boldsymbol{X_{val}}, \boldsymbol{y_{val}}) \leftarrow (\boldsymbol{X_A}, \boldsymbol{y_A})$ // Split the PH pairs
**for** $j$ = 1 **to** $R$ **do** // Run $R$ AL rounds
    **if** j == 1 **do**
        $\boldsymbol{X_s}, \boldsymbol{y_s} \leftarrow sample\_randomly(\boldsymbol{X_{train}}, \boldsymbol{y_{train}})$ // Section 1.1.3.3
        $\boldsymbol{X_r}, \boldsymbol{y_r} \leftarrow (\boldsymbol{X_{train}}, \boldsymbol{y_{train}}) \backslash (\boldsymbol{X_s}, \boldsymbol{y_s})$ // Get the remaining set
    **else do**
        $\boldsymbol{X'_s}, \boldsymbol{y'_s} \leftarrow sample\_actively(model, \boldsymbol{X_r}, \boldsymbol{y_r})$ // Section 1.1.3.3
        $\boldsymbol{X_s}, \boldsymbol{y_s} \leftarrow (\boldsymbol{X_s}, \boldsymbol{y_s}) \cup (\boldsymbol{X'_s}, \boldsymbol{y'_s})$ // Update sampled training set
        $\boldsymbol{X_r}, \boldsymbol{y_r} \leftarrow (\boldsymbol{X_r}, \boldsymbol{y_r}) \backslash (\boldsymbol{X_s}, \boldsymbol{y_s})$ // Get the remaining set
    **end if**
    $Model \leftarrow tuning\_heldout(\boldsymbol{X_s}, \boldsymbol{y_s}, \boldsymbol{X_{val}}, \boldsymbol{y_{val}}, \boldsymbol{H})$ // Select the best model
    $\widehat{\boldsymbol{y_{T_j}}} \leftarrow Model(\boldsymbol{X_T})$
**end for**
$\widehat{\boldsymbol{y_T}} \leftarrow \{\widehat{\boldsymbol{y_{T_1}}}, \widehat{\boldsymbol{y_{T_2}}}, \dots, \widehat{\boldsymbol{y_{T_R}}}\}$
**return** $\{\boldsymbol{X_T}, \widehat{\boldsymbol{y_T}}\}$

**Algorithm 2.** Training the production entailment model and predicting unannotated PH pairs.

**Input:** $\{\boldsymbol{X_A}, \boldsymbol{y_A}, \boldsymbol{X_U}, \boldsymbol{H}\}$, where $\boldsymbol{X_A}$ and $\boldsymbol{y_A}$ are the annotated PH pairs and their labels, $\boldsymbol{X_U}$ is the unannotated PH pairs, and $\boldsymbol{H}$ is the hyperparameter search space.
**Output:** $\{\boldsymbol{X_U}, \widehat{\boldsymbol{y_U}}\}$, where $\boldsymbol{X_U}$ and $\widehat{\boldsymbol{y_U}}$ are the unannotated PH pairs and their corresponding predictions by the production model.

$hparam \leftarrow tuning\_10\_fold(\boldsymbol{X_A}, \boldsymbol{y_A}, \boldsymbol{H})$ // Run grid search

**for** $i = 1$ **to** $S$ **do** // Run $S$ different seeds
  $\boldsymbol{\theta_i} \leftarrow initialize\_random\_weights(hparam)$
  $Model_i \leftarrow train\_model(\boldsymbol{X_A}, \boldsymbol{y_A}, \boldsymbol{\theta_i})$
  $\widehat{\boldsymbol{y}_{\boldsymbol{U}_i}} \leftarrow Model_i(\boldsymbol{X_U})$ // Predict unannotated input
**end for**
$\widehat{\boldsymbol{y}_{\boldsymbol{U}}} \leftarrow get\_elementwise\_mean([\widehat{\boldsymbol{y}_{\boldsymbol{U}_1}}, \widehat{\boldsymbol{y}_{\boldsymbol{U}_2}}, \dots, \widehat{\boldsymbol{y}_{\boldsymbol{U}_S}}])$ // Do ensemble predictions
**Return** $\{\boldsymbol{X_U}, \widehat{\boldsymbol{y}_{\boldsymbol{U}}}\}$

##### 1.2.4 Link prediction

###### 1.2.4.1 Model selection dataset generation

To evaluate different versions of the FAKG in **Fig. 5b** fairly, we created a held-out validation and test set by randomly sampling the triplets of type (food, contains, chemical) from the annotated PH pairs used for the entailment model. Note that we only added the (food, contains, chemical) triplet type, but not the (food part, contains, chemical) triplet type, as we are interested in generating hypotheses of the former. This results in the validation set with 445 positives and the test set with 447 triplets. As for the training data, although the data source varies for each version of the FAKG, the generation process is identical for all. We start by making sure that the 892 (food, contains, chemical) triplets in the validation and test set are removed from the available data. Next, we also remove any (food part, contains, chemical) in the training set that shares the same NCBI taxonomy ID as food entities in the validation and test set. For example, if (strawberry, contains, Ascorbic Acid) is in the validation set, we remove any (strawberry {*part*}, contains, Ascorbic Acid) triplets in the training set. We chose to do this exclusion as we wanted to prevent the model from learning the chemical composition of foods only from their food part chemical composition.

###### 1.2.4.2 Hyperparameter optimization

We performed the hyperparameter optimization of the six link prediction models, TransE[26], ER-MLP[27], DistMult[28], TransD[29], ComplEx[30], and RotatE[31], using the metric MR on the validation set. For each model and each version of the dataset in **Fig. 5b**, we tested 50 different sets of hyperparameters, where each hyperparameter set was drawn randomly from the default pool of hyperparameters of the PyKEEN library[32] that were chosen from the best-reported values in each model's original paper. Early stopping was also used during the hyperparameter optimization process.

###### 1.2.4.3 OpenAI GPT

We tested how the vanilla GPT-3.5 model (text-davinci-003)[33] performs without any training data and how it compares to the graph-embedding models. We used the (food, contains, chemical) triplets in the test set to ask the question "Does {*food*} contains {*chemical*}?" using the text completion endpoint, where the scientific name was used for food, and the PubChem name was used for the chemical (MeSH name was used for chemical in case PubChem entry did not exist). We prompted the model to generate five different versions of the answer to test the statistical significance, similar to what we did with graph-embedding link prediction models. Surprisingly, GPT had a precision of 64.8%, although the recall and F1 score was lower at 31.8% and 42.7%, respectively.

#### 1.3 FoodAtlas Knowledge Graph

##### 1.3.1 Entity types

The current version of FoodAtlas KG (FAKG) supports three entity types: *cellular organism (food)*, *food part*, and *chemical* (**Supplementary Table 4**). All entities in FAKG are assigned a unique FoodAtlas ID, a sequentially assigned numerical ID prefixed with the letter *e*. In this section, we describe each entity type in more detail.

###### 1.3.1.1 *cellular organism* (*food*)

In addition to the unique FoodAtlas ID, *cellular organism* entities have one unique NCBI Taxonomy ID (NCBI ID) and part ID of *p0*, corresponding to the whole food, in contrast to the food parts in the next section. We require all organisms to have a valid NCBI ID to merge external databases and resolve synonyms. Although we interchangeably use the terminology *cellular organism* and *food*, it is worth noting that not all organisms are foods. For example, an entity *strawberry* with a rank *species* in the taxonomic lineage has a parent entity *Fragaria* with a rank *genus* in the knowledge graph. Although both entities are *cellular organism* entities, we call only the entity *strawberry* a *food* entity. Note that in FAKG, this relationship is encoded using a triplet (*Fragaria*, *hasChild*, *strawberry*). While all *food* entities have an outgoing relationship, *contains*, to a *chemical* entity, *cellular organism* entities have no outgoing *contains* relationship but only a *hasChild* relationship to another *cellular organism* or *food* entity.

###### 1.3.1.2 *food part*

Like the *cellular organism* (*food*) entities, each *food part* entity has one unique NCBI ID identical to the corresponding *food* entity (e.g., entities *carrot root* and *carrot* have the same NCBI ID of 4039). In addition, each *food part* entity also has a unique part ID (e.g., *carrot root* has a part ID of *p49*, and *carrot* has a part ID of *p0*). Note that we strictly use the terminology *food part* and not *cellular organism part*, as we derive all *food part* entities from *food* entities, not other *cellular organism* entities.

###### 1.3.1.3 *chemical*

In addition to the unique FoodAtlas ID, each *chemical* entity has either one unique PubChem CID (CID) or MeSH ID. To be specific, a chemical has a unique MeSH ID if we could not find the PubChem ID. We used CID to merge other *chemical* entities from external databases and resolve the synonyms. we prioritized PubChem ID over the MeSH ID as the chemical indexer in FAKG because PubChem is the most comprehensive, used by most external databases, and prevents the dilution of chemicals when merging. The dilution problem happens when multiple identifiers are used to merge the entities. For example, according to PubChem, eight different PubChem entries for *fumaric acid* (CIDs: 6076814, 101823788, 444972, 6364607, 6433510, 6440849, 723, 9793847) all point to a single MeSH entry (MeSH ID: C032005) as a synonym. To make matters worse, each CID can point to more than one MeSH entry or CAS entry. For example, a CID 6076814 has two MeSH entries (MeSH IDs: C032005, C030272) and three CAS entries (CAS IDs: 18016-19-8, 5873-57-4, 7704-73-6). Therefore, if we use more than one unique identifier (PubChem ID in our case) for merging entities and resolving synonyms, we can end up with a chemical entry with multiple chemicals merged (diluted) into one.

##### 1.3.2 Relation types

The current version of FoodAtlas KG supports four relation types: *contains*, *isA*, *hasChild*, and *hasPart* (**Supplementary Table 4**). In addition, all relations in FAKG are assigned a unique FoodAtlas ID, a sequentially assigned numerical ID prefixed with the letter *r*. In this section, we describe each relation type in more detail.

###### 1.3.2.1 *contains*

The *contains* relation type has two possible triplet types (*food*, *contains*, *chemical*) and (*food part*, *contains*, *chemical*), as shown in **Fig. 2a**. These *contains* triplets are either from the FoodAtlas framework or are from three external databases Frida[34], Phenol-Explorer[18–20], and FDC[17].

###### 1.3.2.2 *isA*

The *isA* relation type has one possible triplet type (*chemical*, *isA*, *chemical*) (**Fig. 2a**), which encodes the ontological relationship of the chemical entities. This ontological relationship is taken directly from the MeSH Tree. Note that the head entity is a child node of the tail entity in the MeSH Tree (e.g., (*Catechin*, *isA*, *Flavonoids*)).

###### 1.3.2.3 *hasChild*

The *hasChild* relation type has one possible triplet type (*cellular organism*, *hasChild*, *cellular organism*) (**Fig. 2a**), which encodes the taxonomical relationship of the *cellular organism* entities and is taken directly from the NCBI Taxonomy. In contrast to the *isA* entity type, the tail entity of the *hasChild* triplet is the child node of the head entity (e.g., (*Fragaria*, *hasChild*, *strawberry*)).

###### 1.3.2.4 *hasPart*

The *hasPart* relation type has one possible triplet type (*cellular organism*, *hasPart*, *food part*) (**Fig. 2a**). In the current version of FAKG, the only source of such triplets is the FoodAtlas active learning pipeline. Note that the entailment model only predicts (*food*, *contains*, *chemical*) or (*food part*, *contains*, *chemical*) triplet types, and if a triplet (*food part*, *contains*, *chemical*) is either annotated or predicted as positive by the FoodAtlas pipeline, we automatically extract (*food*, *hasPart*, *food part*) triplet from it and mark it as a positive.

##### 1.3.3 Knowledge injection

We first inject *contains* and *hasPart* triplets into the FoodAtlas KG, followed by *isA* triplets and *hasChild* triplets from the MeSH Tree and NCBI Taxonomy, respectively. This section describes the exact procedures for this knowledge injection process.

###### 1.3.3.1 FoodAtlas active learning pipeline

Three triplet types, (*food*, *contains*, *chemical*), (*food part*, *contains*, *chemical*), and (*food*, *hasPart*, *food part*), are injected into FAKG from the FoodAtlas active learning (AL) pipeline. The *food* and *chemical* entities are originally tagged with NCBI IDs and MeSH IDs, respectively, by the LitSense API. To inject these triplets into the FoodAtlas KG, a preprocessing step of looking up the corresponding PubChem CIDs (CIDs) for the *chemical* entities is necessary, as CID is used to merge the entities (**Section 1.3.1.3 of Supplementary Information**). For example, a *chemical* entity of *oleic acid* from the triplet (*food*, *contains*, *oleic acid*) has a unique MeSH ID of D019301. We search this unique MeSH ID in the PubChem database and find all PubChem compound entries that list the

MeSH ID as a synonym. Then, for *oleic acid*, three CIDs (445639, 5460221, 965) all list the MeSH ID of D019301 as a synonym. Therefore, the original triplet (*food*, *contains*, *oleic acid*) is expanded into three triplets, each with a unique CID, before being injected into the FoodAtlas KG.

###### 1.3.3.2 External databases

We inject the (*food*, *contains*, *chemical*) triplets from three external DBs, Frida, Phenol-Explorer, and FDC, into FAKG. We describe the exact steps for processing data from these external DBs in detail in **Section 1.3.4 of Supplementary Information**. The *food* entities from these external DBs are already tagged with NCBI taxonomy, whereas the *chemical* entities are tagged with either CID or CAS ID but not MeSH ID. If a *chemical* entity has a CID, we use it to search the PubChem database for synonym MeSH IDs. We then associate the found MeSH IDs to the CID, which are further used to merge the entities as described in **Section 1.3.1.3 of Supplementary Information**. However, if a *chemical* entity does not have a CID but only has a CAS ID, we look up this CAS ID on PubChem and, in turn, retrieve the MeSH IDs. Like MeSH, a single CAS ID can be a synonym for multiple PubChem entries. In such cases, we perform the same process of expanding the original triplet to multiple triplets, each with a unique CID. Note that chemicals added from external DBs may not always have a MeSH ID since some PubChem entries do not have a related MeSH ID.

###### 1.3.3.3 MeSH

Once all *chemical* entities are added to FAKG, we add the ontological relationships of these chemicals using the MeSH Tree. For all *chemical* entities in the KG, we find the subset of entities with a unique MeSH ID (19,645 out of 19,722) and then search the MeSH database for these MeSH IDs. There are two types of MeSH entries: *MeSH descriptor data*, which has a prefix of the letter *D* in its unique ID (e.g., D002392 for catechin), and *MeSH supplementary concept data*, which has a prefix of the letter *C* in its unique ID (e.g., C024603 for Geraniin). All descriptor data with the prefix *D* has one or more MeSH Tree numbers. For example, catechin (MeSH ID: D002392) has 4 MeSH Tree numbers, as shown in **Supplementary Fig. 9a**. Each MeSH Tree is converted to a triplet by connecting the tree's nodes with the *isA* relation type. For example, catechin with MeSH Tree number D03.383.663.283.240.190 in **Supplementary Fig. 9a** will be expanded into five triplets (*Catechin*, *isA*, *Chromans*), (*Chromans*, *isA*, *Benzopyrans*), (*Benzopyrans*, *isA*, *Pyrans*), (*Pyrans*, *isA*, *"Heterocyclic Compound, 1-Ring"*), (*"Heterocyclic Compounds, 1-Ring"*, *isA*, *Heterocyclic Compounds*). All supplementary concept data with the prefix *C* are mapped to one or more descriptor data with the prefix *D*. Geraniin (MeSH ID: C024603), for example, is mapped to two descriptor data Glucosides (MeSH ID: D005960) and Hydrolyzable Tannins (MeSH ID: D047348). We encode this relationship with a triplet (*Geraniin*, *isA*, *Glucosides*) and (*Geraniin*, *isA*, *"Hydrolyzable Tannins"*), respectively. The MeSH tree of the two descriptor entities Geraniin mapped to is also translated into triplets and injected into FAKG.

###### 1.3.3.4 NCBI Taxonomy

For all *food* entities in the KG, their corresponding NCBI Taxonomy IDs are used to search the NCBI Taxonomy to retrieve the taxonomic lineage information. For example, the complete lineage data of strawberry (NCBI:txid3747) with rank *species* are shown in

**Supplementary Fig. 10**, where the cellular organism (NCBI:txid131567) is the root entity of the lineage, and the Fragaria (NCBI:txid3746) is the immediate parent entity of rank *genus*. This lineage is converted to triplets by connecting the entities in the lineage with *hasChild* relationship as (*cellular organism*, *hasChild*, *Eukaryota*), (*Eukaryota*, *hasChild*, *Viridiplantae*), …, and (*Fragaria*, *hasChild*, *Fragaria x ananassa*).

##### 1.3.4 Triplets from external databases

This section will discuss how we integrate triplets extracted from Frida, FDC, and Phenol-Explorer. Note that we have considered other food chemical composition databases. However, to integrate with FAKG, the databases need to have standardized food and chemical indexers consistent with FAKG; Specifically, a food must have an NCBI ID, and a chemical must have a CID. Most databases, however, do not necessarily suffice this requirement; For example, Frida does not provide food NCBI IDs, and FDC does not provide CIDs. In the next paragraph, we will discuss how to rely on our metadata retrieval pipeline to retrieve the NCBI IDs if the database provides scientific names. However, if a database lacks CIDs, we need to contact the authors of the databases to receive chemical IDs that can be linked to CIDs internally. The databases integrated into FAKG either contain or have provided CIDs or IDs linked to CIDs to us internally. We are actively working with the authors of other databases, including FooDB, to expand FAKG in the following versions.

To integrate triplets from FDC, Frida, and Phenol-Explorer into FAKG with a consistent standard, we have developed a semi-automatic method relying on NCBI Entrez[35] to retrieve NCBI IDs, MeSH IDs, and PMIDs with scientific names, other chemical IDs, and publication titles, respectively (**Section 1.3.4.1 of Supplementary Information**).

Another property of FoodAtlas is its focus on raw foods: Food products (e.g., chocolate) or processed foods (e.g., roasted chicken) may contain inconsistent chemical additives, which introduces noise to FAKG. Thus, we relied on a rule-based filterer to remove the food-chemical relations that were non-raw foods for each external source (**Section 1.3.4.2 of Supplementary Information**).

For the rest of this section, we will discuss the general methodology of metadata retrieval and relation filtering pipelines in the first two subsections. Then, we will summarize the specific treatment and the result for each external database.

###### 1.3.4.1 Metadata Retrieval

**Retrieving NCBI IDs**. If the scientific name of a food is available, we may use NCBI Entrez to query the NCBI ID with the following steps:

- Make the scientific name lowercase.
- Report for manual validation if the input contains the “x” or “u’\xd7’” term.
    - Note: Crossbreeds are hard to parse but rare, so they are dealt with manually.
- Drop the input with less than two terms.
    - Note: It cannot be species, subspecies, or varieties.
- Drop the input with “.” in the first two terms.
    - Note: Scientific names for species, subspecies, or varieties have no abbreviations in the first two terms.

- Report for manual validation if the input contains “convar” or “convar.”.
  - Note: Convarieties are hard to parse but rare, so they are dealt with manually.
- Report for manual validation if the input contains more than two terms and does not contain “.”.
  - Note: The majority of the scientific names have lengths of 2. If more than 2, the scientific names either contain the abbreviated authority, e.g., “L.”, or variety terms, such as “subsp.” and “var.”. NCBI Entrez does strict string matching for NCBI ID queries, so this step ensures that data sources are not omitting “.” by accident.
- If the scientific name has a length of 2, query the NCBI ID.
- If the scientific name has a length longer than 2, format the variety terms, remove the authority terms, and query the NCBI ID.
  - Note: Some external databases use the variety terms that are not accepted by NCBI Entrez, e.g., “ssp.” instead of “subsp.”, which needs to be formatted accordingly. The authority terms are optional for the NCBI Entrez API. However, if authority terms are contained, they must be correct for the API to return the NCBITaxon ID. Thus, it is much easier to remove them.
- Report for manual validation if NCBI Entrez does not return an ID. Fix if the error is due to minor typos.

This procedure ensured that the automated part of our method was exact (i.e., if NCBI Entrez returned, it was correct) and minimized the need for manual validation.

**Retrieving PMID**. If the publication title of a reference is available, we may use NCBI Entrez to query the PMID with the following steps:

- Make the publication title lowercase.
- Remove all stop words and reformat the input into the advanced search query format defined in NCBI Entrez API[35].
  - Note: Specifically, the underlined terms of “uptake and metabolism of epicatechin and its access to the brain after oral ingestion” are removed first, and then the sentence is transformed to “uptake[Title] AND metabolism[Title] AND epicatechin[Title] AND access[Title] AND brain[Title] AND after[Title] AND oral[Title] AND ingestion[Title]”. We have empirically tested and found that this approach maximizes the hit rate.
- Query with the formatted input and retrieve its PMID(s).
- Select and verify the correctness of the PMID with the following steps.
  - Use the retrieved PMID to query and retrieve the publication title from PubMed.
  - Make the retrieved publication title lowercase.
  - Remove all punctuations and whitespaces in the input publication title and the retrieved publication title.
  - Compare the input publication title and the retrieved publication title. Verification succeeds if and only if the two titles are strictly matching. If not, report for manual validation.

- If multiple PMIDs are returned for a single input publication title, report for manual validation if none of the PMIDs pass the verification.

Similar to retrieving NCBITaxon ID, this procedure ensured that our method's automated part was exact (i.e., if it passed verification, it was correct) while minimizing manual validation.

###### 1.3.4.2 Relation Filtering

Due to the inconsistency of food naming conventions among external databases, relation filtering has been implemented specifically for each external database to remove relations associated with non-raw foods. The primary mechanism of the filtering is to detect either specific substring of food names or specific food groups if available. If the concentration value associated with a relation is zero, we do not add that relation to the KG.

###### 1.3.4.3 FDC

**Foods**. FDC is a massive reservoir of food. However, after consulting with researchers working on FDC, since the SR Legacy foods samples mixed different species/subspecies, we decided not to use them. Therefore, we only focused on Foundation Foods, where 56 have NCBI IDs.

**Chemicals**. As of 02/03, the current database does not provide CIDs or CAS IDs for the chemicals in the database. We consulted the FDC researchers and received some work-in-progress ID mapping internally. With that, we retrieved CIDs for 62 chemicals.

###### 1.3.4.4 Frida

**Foods**. Frida contains 1,249 unique foods with common and scientific names. To exclude foods that are food products or processed (i.e., with additives), we only included foods with the term *raw* in their food names, excluding most non-raw foods. In addition, we ignored some food groups (**Supplementary Table 4**) for further food cleaning. Lastly, we retrieved NCBI IDs for these foods using their scientific names, which resulted in 200 usable raw foods.

**Chemicals**. Frida contains 205 unique *parameters*, where a parameter can be either a micronutrient or a macronutrient. We dropped most macronutrients due to their lacking CIDs. Because the original Frida database does not contain CIDs, we contacted the correspondent of the Frida team and internally received the CIDs and CAS IDs for its parameters. Finally, we collected 133 chemicals compatible with FAKG.

###### 1.3.4.5 Phenol-Explorer

**Foods**. Phenol-Explorer initially contains 459 foods with common and scientific names. After removing products and processed foods (**Supplementary Table 4**), we retrieved NCBI IDs using the metadata retrieval pipeline. Consequently, we obtained 194 foods with IDs.

**Chemicals**. Phenol-Explorer contains 501 chemicals, each, if available, associated with CID and CAS ID. We used 155 chemicals that were at least associated with one of the IDs.

### 2 Supplementary Figures

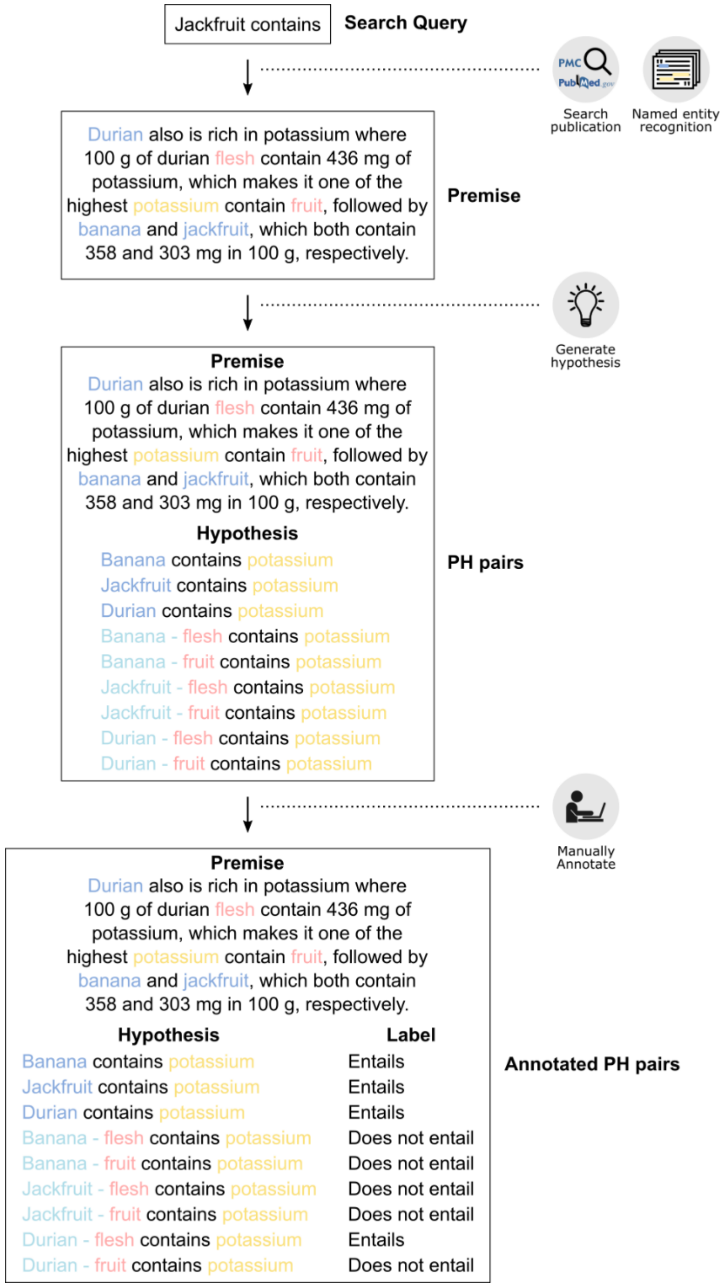

**Supplementary Fig. 1.** Example of PH pairs generation and the annotation process of the FoodAtlas framework. For a single premise shown above, 9 PH pairs are generated, among which four are annotated as being positives.

a

| PH pair | Probability |
|---|---|
| #1 | 0.99 |
| #2 | 0.95 |
| #3 | 0.86 |
| #4 | 0.83 |
| #5 | 0.72 |
| #6 | 0.69 |
| #7 | 0.66 |
| #8 | 0.50 |
| #9 | 0.42 |
| #10 | 0.39 |
| #11 | 0.25 |
| #12 | 0.22 |
| #13 | 0.15 |
| #14 | 0.09 |
| #15 | 0.02 |

b

| PH pair | Probability |
|---|---|
| #1 | 0.99 |
| **#2** | **0.95** |
| #3 | 0.86 |
| #4 | 0.83 |
| #5 | 0.72 |
| #6 | 0.69 |
| #7 | 0.66 |
| #8 | 0.50 |
| **#9** | **0.42** |
| #10 | 0.39 |
| #11 | 0.25 |
| #12 | 0.22 |
| **#13** | **0.15** |
| #14 | 0.09 |
| #15 | 0.02 |

c

| PH pair | Probability |
|---|---|
| **#1** | **0.99** |
| **#2** | **0.95** |
| **#3** | **0.86** |
| #4 | 0.83 |
| #5 | 0.72 |
| #6 | 0.69 |
| #7 | 0.66 |
| #8 | 0.50 |
| #9 | 0.42 |
| #10 | 0.39 |
| #11 | 0.25 |
| #12 | 0.22 |
| #13 | 0.15 |
| #14 | 0.09 |
| #15 | 0.02 |

d

| PH pair | Probability | Uncertainty |
|---|---|---|
| #1 | 0.99 | 0.01 |
| #2 | 0.95 | 0.05 |
| #3 | 0.86 | 0.14 |
| #4 | 0.83 | 0.17 |
| #5 | 0.72 | 0.28 |
| #6 | 0.69 | 0.31 |
| **#7** | **0.66** | **0.34** |
| **#8** | **0.50** | **0.50** |
| **#9** | **0.42** | **0.42** |
| #10 | 0.39 | 0.39 |
| #11 | 0.25 | 0.25 |
| #12 | 0.22 | 0.22 |
| #13 | 0.15 | 0.15 |
| #14 | 0.09 | 0.09 |
| #15 | 0.02 | 0.02 |

e

| PH pair | Probability |
|---|---|
| #1 | 0.99 |
| **#2** | **0.95** |
| #3 | 0.86 |
| #4 | 0.83 |
| #5 | 0.72 |
| #6 | 0.69 |
| #7 | 0.66 |
| #8 | 0.50 |
| #9 | 0.42 |
| **#10** | **0.39** |
| #11 | 0.25 |
| **#12** | **0.22** |
| #13 | 0.15 |
| #14 | 0.09 |
| #15 | 0.02 |

**Supplementary Fig. 2.** Visualization of the active learning sampling strategies where three PH pairs (marked in bold) are to be chosen for each strategy. **a**, A sample of 15 PH pairs for visualization purposes was ordered from high probability to low. **b**, The *stratified* sampling strategy first bins the PH pairs into three equally sized bins, and one sample is drawn from each bin. Note that in actual implementation, we bin the PH pairs into ten equal-sized bins. **c**, The *maximum likelihood* strategy selects the top three PH pairs with the highest probability. **d**, The *maximum entropy* strategy selects the PH pairs with the highest uncertainty scores. **e**, The *random* strategy selects PH pairs randomly.

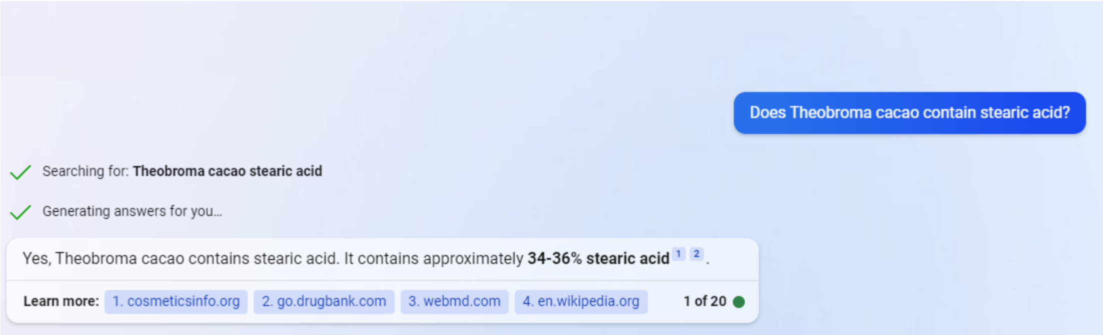

**Supplementary Fig. 3.** Sample illustration of a Bing Chat search. In this example, Bing Chat returned two references to its claims. We made decisions by checking the validity of the referenced sources. Note that the conversation type of Bing Chat was set to *Balanced*.

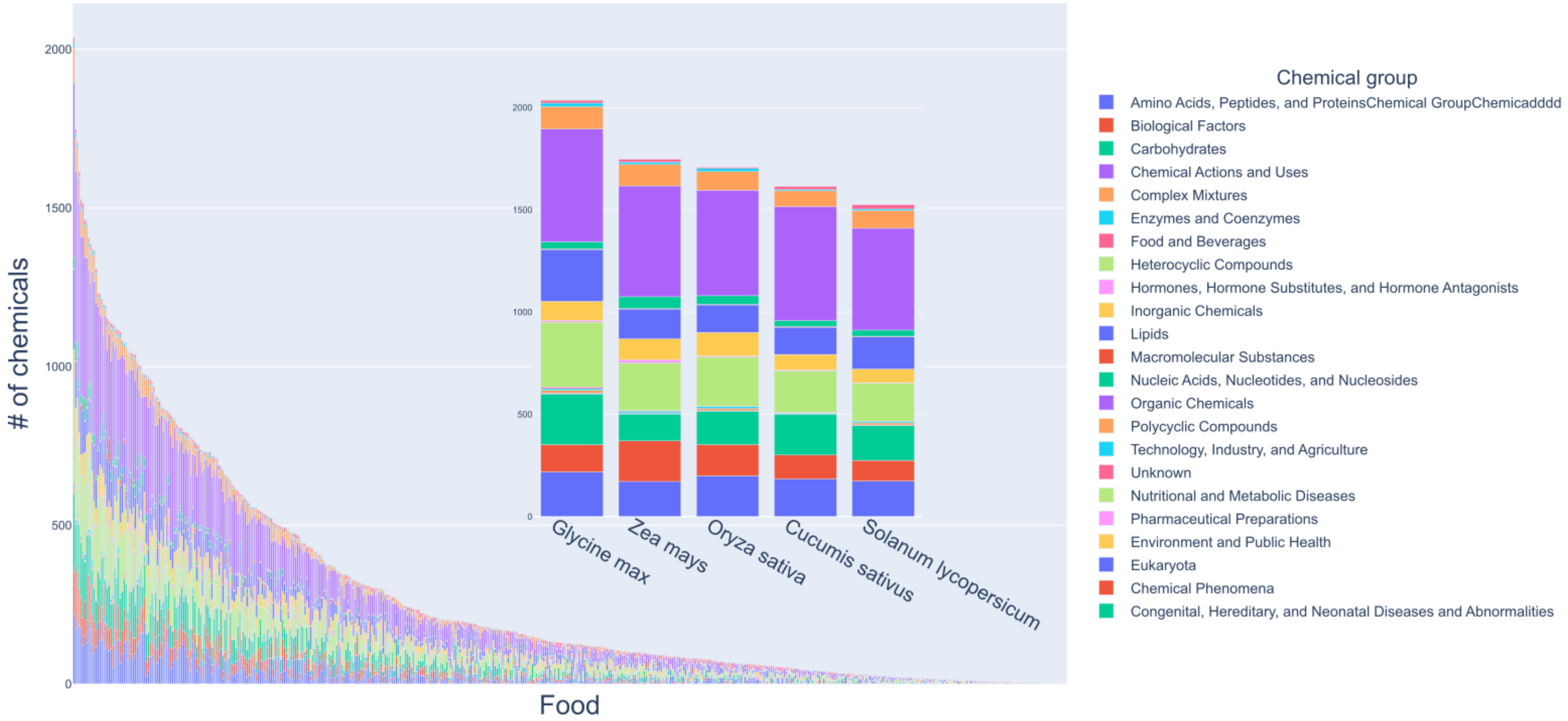

**Supplementary Fig. 4**. Foods in the FoodAtlas knowledge graph and their number of connections to the chemicals by the *contains* relationship. The chemicals are color-coded by their chemical group. The barplot inside the main bar plot shows the top 5 foods with the most *contains* relationship in the knowledge graph.

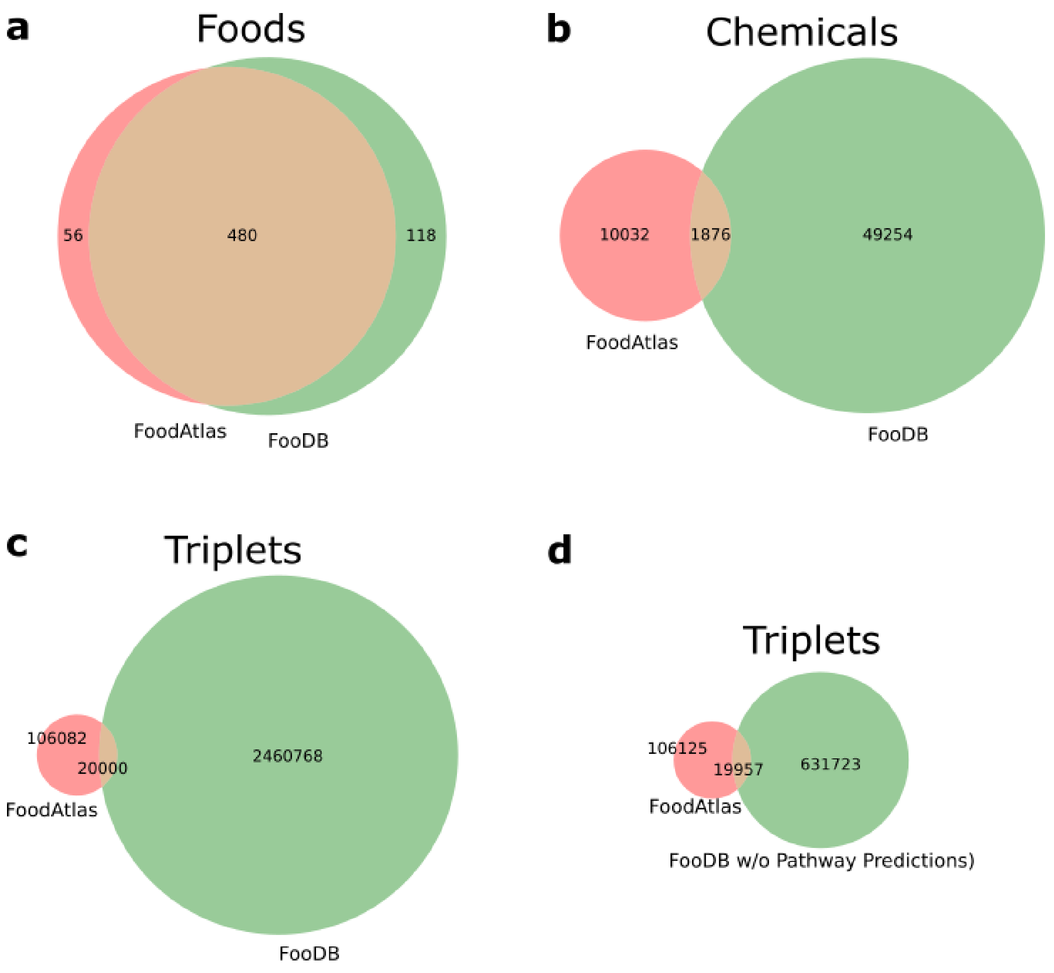

**Supplementary Fig. 5.** Comparison of the number of entities in FAKG with that in FooDB. **a**, **b**, Foods and chemicals were indexed based on NCBI Taxonomy IDs and PubChem CIDs, respectively, and the Venn diagrams show the coverage of unique IDs of the two databases. **c**, **d**, Triplets were indexed based on pairs of NCBI Taxonomy IDs and PubChem CIDs, and the diagrams show the coverage of the unique paired IDs of the two databases. Note that a food with food part has the same NCBI Taxonomy ID as the corresponding food, and thus it was considered as one entity in the diagrams. See **Supplementary Information Section 1.1.9** for more information.

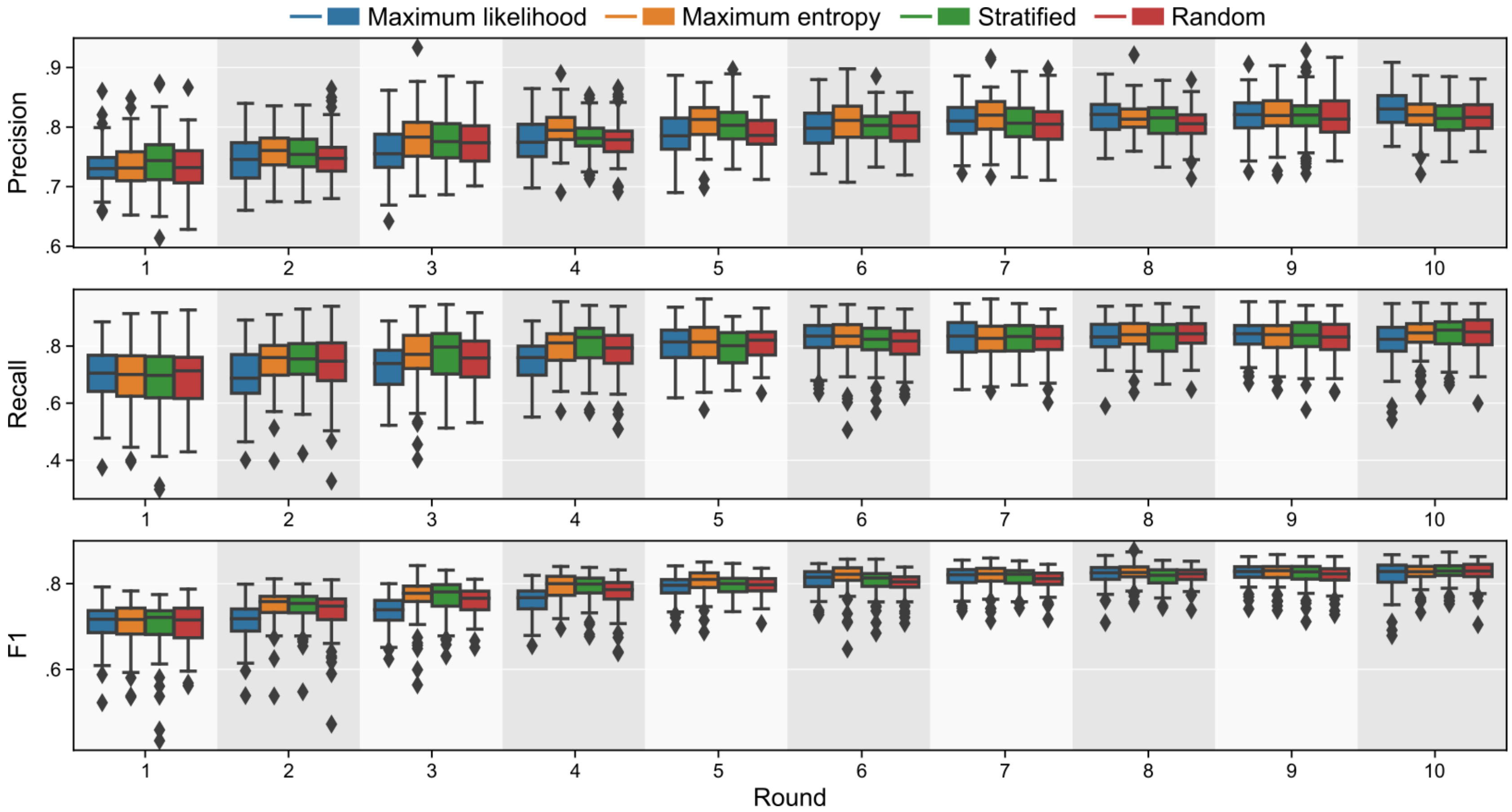

**Supplementary Fig. 6.** Precision, recall, and F1 score of four AL sampling methods over the ten rounds of AL. The box represents the interquartile range, the middle line represents the median, the whisker line extends from minimum to maximum values, and the diamond represents outliers.

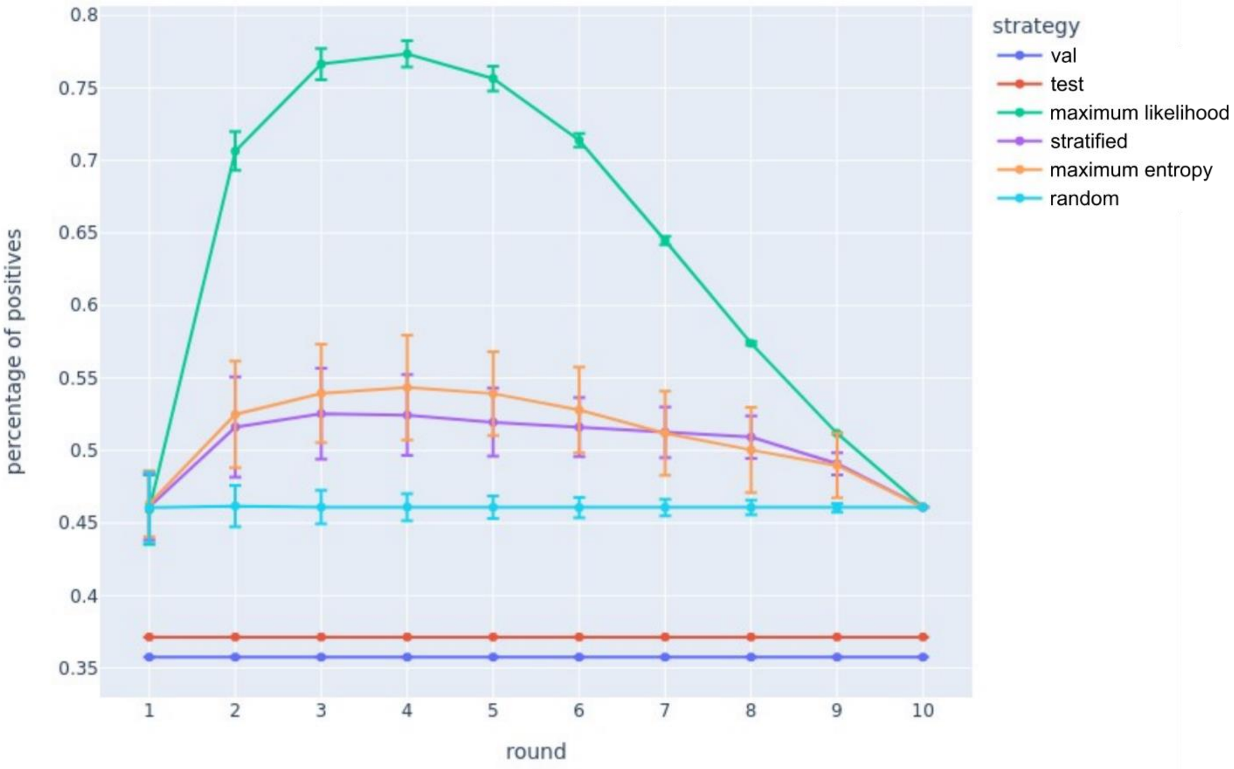

**Supplementary Fig. 7.** Percentage of the positive PH pairs in the training data for each round of 4 different active learning strategies, validation set, and test set. The error bars represent the standard deviation of the percentage of positives (*n* = 100 for 100 random seeds).

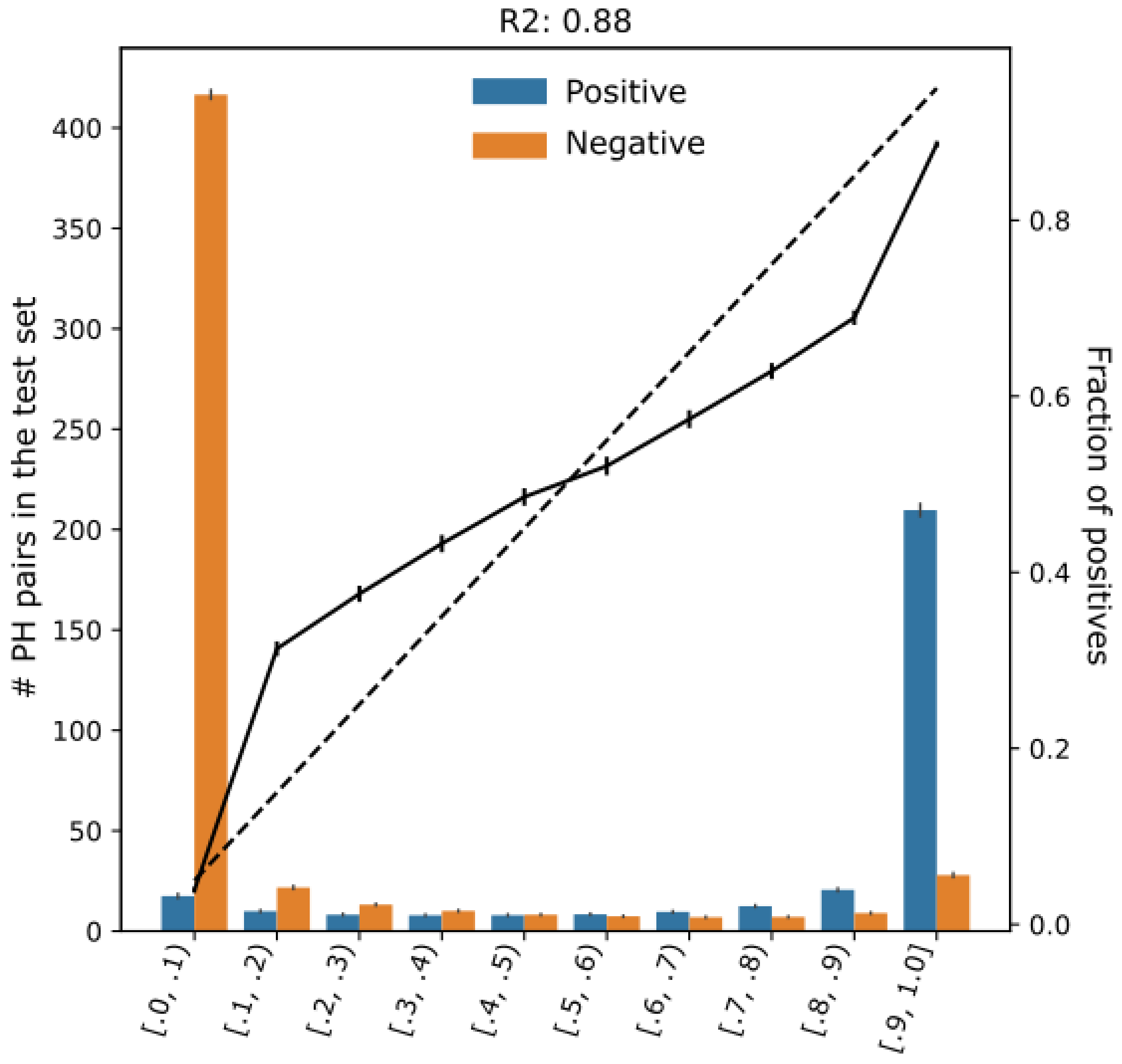

**Supplementary Fig. 8.** Calibration plot of the entailment model with ten bins. The 5-bin $R^2$ was lower than that of the 10-bin (0.88 vs. 0.94), which was expected due to the small number of samples in the intermediate bins.

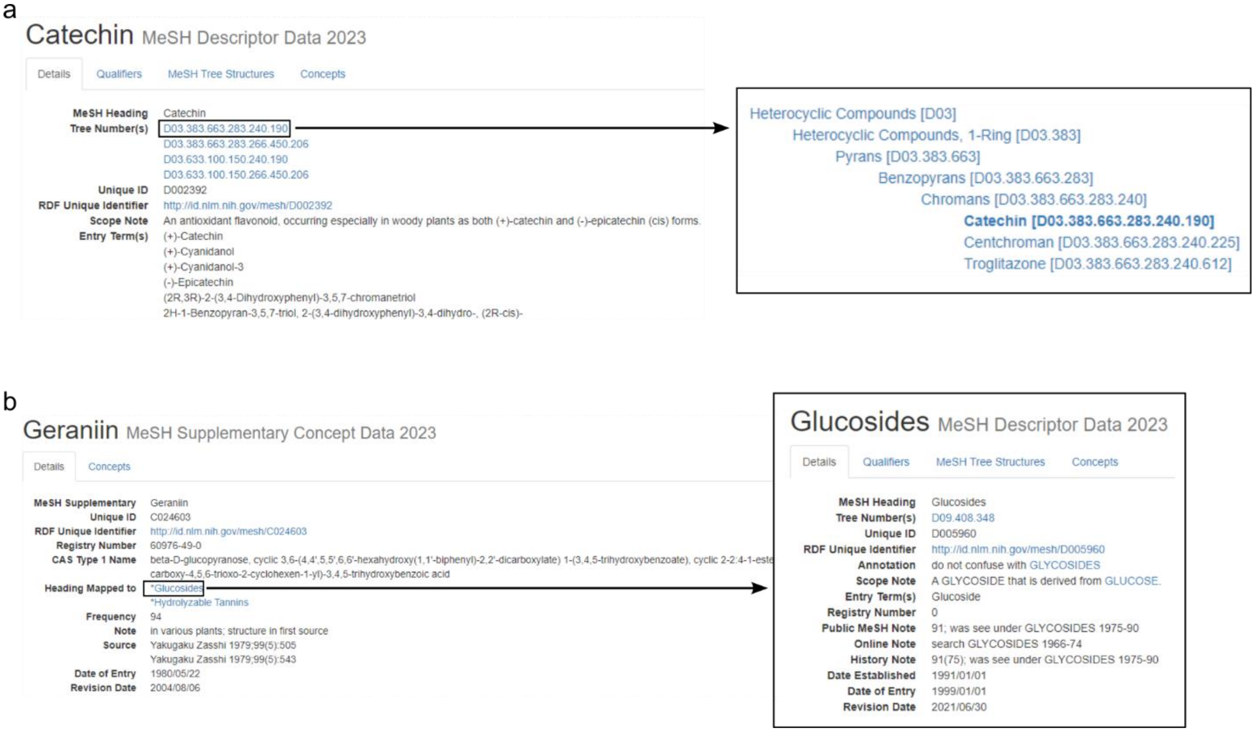

**Supplementary Fig. 9.** Sample illustration of querying two types of MeSH data. **a**, Catechin is a MeSH descriptor data that has a unique MeSH ID with the prefix 'D'. All descriptor data have one unique MeSH ID and can have one or more MeSH tree numbers (4 in the case of catechin). Each tree number encodes the ontological relationships of that MeSH entry in the hierarchical format as seen on the right. **b**, Geraniin is MeSH supplementary concept data that has a unique MeSH ID with the prefix 'C'. All supplementary concept data have one unique MeSH ID, but they do not have a MeSH tree number. Instead, supplementary concept data is mapped to one or more descriptor data (e.g., Geraniin is mapped to Glucosides and Hydrolyzable Tannins).

#### Fragaria x ananassa

Taxonomy ID: 3747 (for references in articles please use NCBI:txid3747)

current name

***Fragaria* × *ananassa*** (Weston) Duchesne ex Rozier

Genbank common name: **strawberry**
NCBI BLAST name: **eudicots**
Rank: **species**
Genetic code: Translation table 1 (Standard)
Mitochondrial genetic code: Translation table 1 (Standard)
Plastid genetic code: Translation table 11 (Bacterial, Archaeal and Plant Plastid)
Other names:

heterotypic synonym

***Fragaria ananassa***

heterotypic synonym

***Fragaria chiloensis* × *Fragaria virginiana***

heterotypic synonym

***Fragaria virginiana* × *Fragaria chiloensis***

Lineage*( full )*
cellular organisms; Eukaryota; Viridiplantae; Streptophyta; Streptophytina; Embryophyta; Tracheophyta; Euphyllophyta; Spermatophyta; Magnoliopsida; Mesangiospermae; eudicotyledons; Gunneridae; Pentapetalae; rosids; fabids; Rosales; Rosaceae; Rosoideae; Potentilleae; Fragariinae; Fragaria

**Supplementary Fig. 10.** Sample illustration of an NCBI taxonomy entry. Strawberry (Fragaria x ananassa) with NCBI taxonomy ID 3747 has a taxonomic lineage with a root node ‘cellular organism’ down to an immediate parent node ‘Fragaria’.

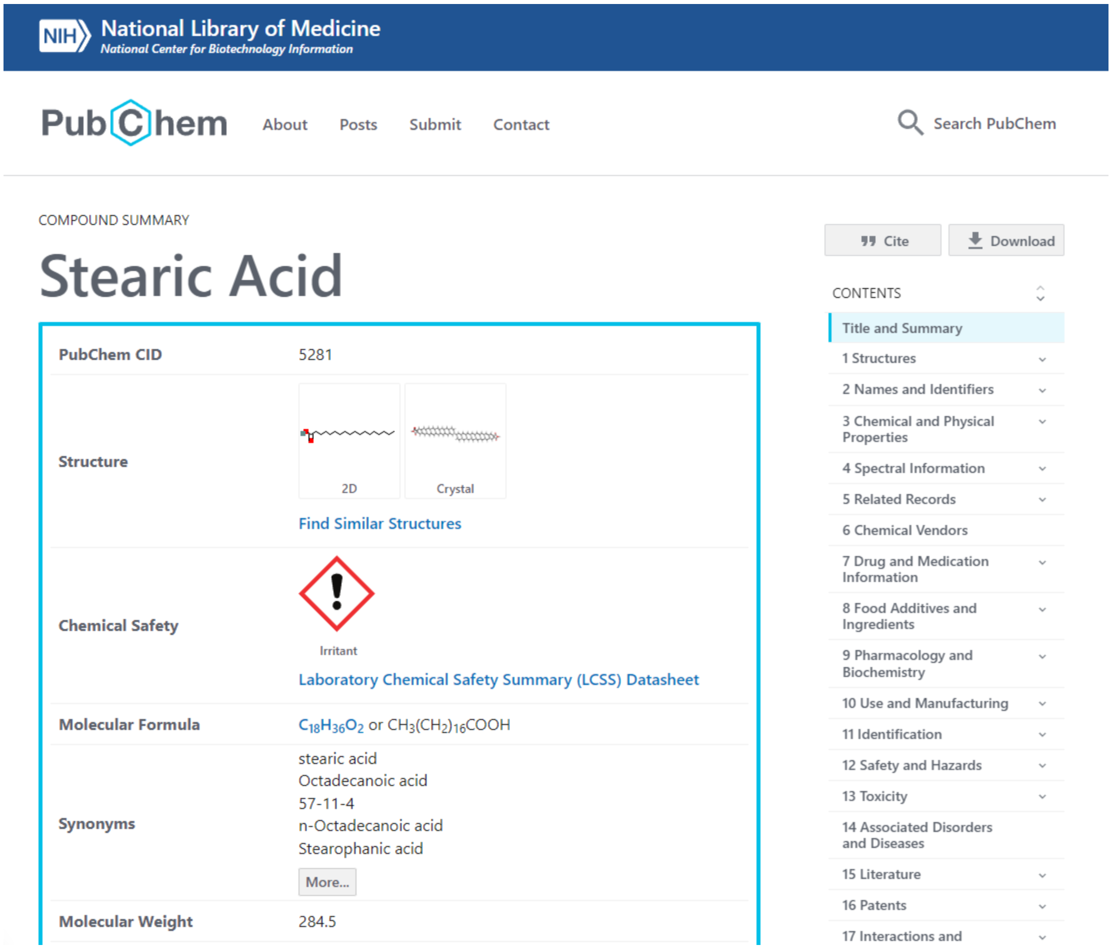

**Supplementary Fig. 11.** Illustration of a PubChem entry. *Stearic acid* is the most common chemical name given its structure. When the most common name did not return evidence during the link prediction validation query, we used the non-ID synonyms shown on the front page. In this example, the synonyms used for searching were *Octadecanoic acid*, *n-Octadecanoic acid*, and *Stearophanic acid*. PubChem heuristically sorts chemical names based on [https://pubchem.ncbi.nlm.nih.gov/docs/compounds#section=Name-Weighting](https://pubchem.ncbi.nlm.nih.gov/docs/compounds#section=Name-Weighting). There are more synonyms if the user clicks the *More* icon, but we decided not to use the complete list of synonyms, which would excessively increase the search space.

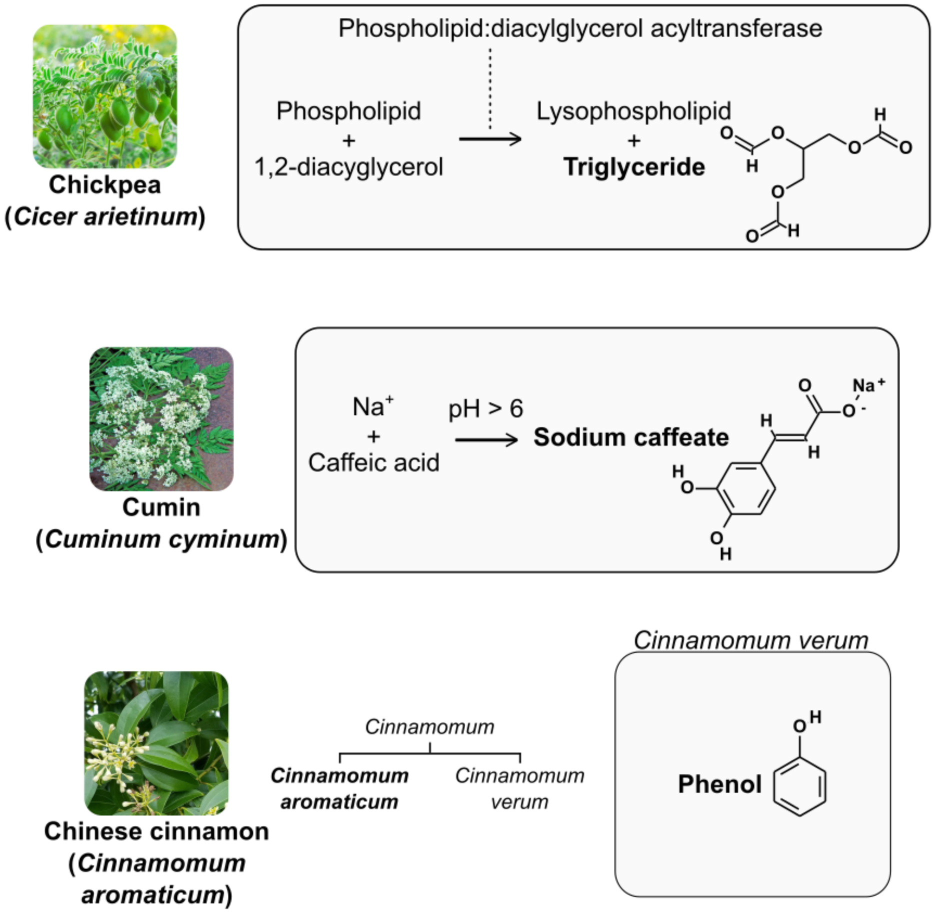

**Supplementary Figure 12.** Three additional food-chemical relationships are not directly mentioned in the literature but with indirect evidence. The possible reason triglyceride and phenol relationships are not mentioned is that the literature rarely mentions them as specific chemical structures, e.g., triglycerides and phenols mean chemical groups that triglyceride (CID: 5460048) and phenol (CID: 996) belong, respectively. Similarly, caffeic acid is commonly found in plants, where it naturally forms sodium caffeate, which can result in the relationship being too obvious to be reported in the literature. Nonetheless, these relationships served as positive controls, such that our link prediction worked as intended.

### 3 Supplementary Tables

**Supplementary Table 1.** Confusion matrix of the entailment models (400 models from four active learning strategies) trained on the entire training dataset (the final active learning round) for the test set.

| | | **Ground Truth** | | | | |
|---|---|---|---|---|---|---|
| | | **Positive** | **Negative** | | | |
| **Prediction** | **Positive** | 260.6 ± 20.3 | 58.3 ± 15.3 | 318.9 ± 33.9 | 0.82 ± 0.03 | **Precision** |
| | **Negative** | 51.4 ± 20.3 | 469.7 ± 15.3 | 521.1 ± 33.9 | 0.90 ± 0.03 | **NPV** |
| | | 312 | 528 | | | |
| | | 0.84 ± 0.07 | 0.89 ± 0.03 | 0.87 ± 0.01 | 0.83 ± 0.03 | |
| | | **Recall** | **Specificity** | **Accuracy** | **F1** | |

**Supplementary Table 2.** Confusion matrix of the PH pairs in the test set separated by which section the premise is from. In addition to the named entity recognition results for each premise, LitSense API also returns where in the literature the premise is extracted from. N/A refers to any other sections aside from the commonly used 8 sections listed below.

| Section | Precision | Recall | F1 | TP | FP | FN | TN |
|---|---|---|---|---|---|---|---|
| N/A | 0.84 | 0.93 | 0.88 | 468 | 86 | 37 | 499 |
| Intro | 0.91 | 0.94 | 0.93 | 602 | 57 | 40 | 206 |
| Discussion | 0.83 | 0.93 | 0.88 | 285 | 57 | 20 | 160 |
| Abstract | 0.77 | 0.88 | 0.82 | 484 | 144 | 66 | 1,000 |
| Methods | 0.89 | 0.81 | 0.85 | 81 | 10 | 19 | 613 |
| Results | 0.80 | 0.88 | 0.84 | 254 | 63 | 36 | 194 |
| Conclusion | 0.74 | 0.97 | 0.84 | 35 | 12 | 1 | 31 |
| Title | 0.75 | 0.89 | 0.81 | 65 | 22 | 8 | 89 |
| Table | 0.33 | 1.00 | 0.50 | 1 | 2 | 0 | 31 |

**Supplementary Table 3.** Potential food-chemical associations generated by the link-prediction (LP) model. Validation of LP-generated (food, contains, chemical) hypotheses with scientific literature identified 355 of them to be true. However, we did not find any evidence for the following 11 pairs with a probability >80%.

| **Food (scientific name; NCBI taxonomy)** | **Chemical (PubChem CID)** | **Probability** |
|---|---|---|
| Japanese persimmon (*Diospyros kaki*; 35925) | 3-Rhamnosyl-Glucosyl Quercetin (156963207) | 92.0% ± 2.9% |
| Chervil (*Anthriscus cerefolium*; 40888) | Loxoprofen (3965) | 91.9% ± 3.0% |
| Dudaim melon (*Cucumis melo var. dudaim*; 2034236) | Matairesinol (119205) | 90.5% ± 6.4% |
| Bearded tooth (*Hericium erinaceus*; 91752) | Lumisterol (6436872) | 90.2% ± 6.2% |
| Chive (*Allium schoenoprasum*; 74900) | Salicylic acid (338) | 90.1% ± 5.9% |
| Chickpea (*Cicer arietinum*; 3827) | Triglyceride (5460048) | 90.1% ± 3.1% |
| Cumin (*Cuminum cyminum*; 52462) | Sodium caffeate (23694762) | 90.0% ± 3.2% |
| Chinese cinnamon (*Cinnamomum aromaticum*; 119260) | Phenol (996) | 87.0% ± 5.9% |
| Atlantic cod (*Gadus morhua*; 8049) | Beta-carotene (5280489) | 84.3% ± 9.2% |
| Butternut (*Juglans cinerea*; 91214) | 2-Hydroxy-1,4-naphthoquinone (6755) | 83.4% ± 13.9% |

| | | |
|---|---|---|
| Turmeric (*Curcuma longa*; 136217) | 3-Rhamnosyl-Glucosyl Quercetin (156963207) | 81.1% ± 10.4% |

**Supplementary Table 4.** Different sources of data in the FoodAtlas KG and their corresponding triplet types.

| Source | Head Type | Relation Type | Tail Type | Quality |
|---|---|---|---|---|
| $FoodAtlas_{annotation}$ | *Cellular organism (food)* | *contains* | *chemical* | High |
| $FoodAtlas_{annotation}$ | *food - part* | *contains* | *chemical* | High |
| $FoodAtlas_{annotation}$ | *Cellular organism (food)* | *hasPart* | *food - part* | High |
| $FoodAtlas_{entailment\_prediction}$ | *Cellular organism (food)* | *contains* | *chemical* | Medium |
| $FoodAtlas_{entailment\_prediction}$ | *food - part* | *contains* | *chemical* | Medium |
| $FoodAtlas_{entailment\_prediction}$ | *Cellular organism (food)* | *hasPart* | *food - part* | Medium |
| $FoodAtlas_{link\_prediction}$ | *Cellular organism (food)* | *Contains* | *Chemical* | Low |
| $FoodAtlas_{MeSH}$ | *chemical* | *isA* | *chemical* | Medium |
| $FoodAtlas_{NCBI}$ | *Cellular organism* | *hasChild* | *Cellular organism* | Medium |
| $FoodAtlas_{Frida}$ | *Cellular organism (food)* | *contains* | *chemical* | Medium & Low |
| $FoodAtlas_{Phenol-Explorer}$ | *Cellular organism (food)* | *contains* | *chemical* | Medium & Low |
| $FoodAtlas_{FDC}$ | *Cellular organism (food)* | *contains* | *chemical* | Low |

**Supplementary Table 5.** Food groups excluded in external databases.

| Frida | Phenol-Explorer |
|---|---|
| Biscuits and cookies | Alcoholic beverages |
| Boiled, smoked, cured or dried meat | Coffee and cocoa |
| Breast milk and infant formula | Cereal products |
| Canned fruit products | Cocoa beverage – Chocolate |
| Canned legumes | Coffee beverage - Arabica<br>Coffee beverages |
| Canned vegetable products | Coffee beverage - Robusta<br>Coffee beverages |
| Cold cuts | Coffee beverage - Unknown<br>Coffee beverages |
| Condiments | Jams - Berry jams |
| Fermented milk products | Jams - Drupe jams |
| Firm rennet cheese | Jams - Pome jams |
| Marmelade, jelly etc. | Other seasonings |
| Other legume products | Soy and soy products |
| Other meat and fresh meat products | Soy drinks |
| Other vegetable products | Spices - Spice blends |
| Potato chip and snacks | Tea infusions |
| Processed cheese | |
| Unfermented milk products | |
| Yeast and baking powder | |

**Supplementary Table 6.** Comparison of the number of associations in external databases. We considered a reference indexed if the corresponding ISSN or journal title could be found in at least one of AGRICOLA, CABI, WoS, and Scopus. Food was considered indexed if it was associated with any identifier (including scientific name). A chemical was considered indexed if it was associated with any identifier, not including chemical formula or molecular weight. Note that the numbers of the final column are different from the numbers of triplets merged into FoodAtlas because FoodAtlas requires triplets to not only be indexed but also to specifically have NCBI Taxonomy IDs for foods and PubChem CIDs or MeSH IDs for chemicals; Also, triplets without evidence, e.g., FDC, were added as low-quality triplets to FoodAtlas.

| Source | # Associations | # Associations w/ ref. (# Ref.) | # Associations w/ indexed ref. (# Indexed ref.) | # Associations w/ indexed ref., food, and chemical |
|---|---|---|---|---|
| FDC | 135,073 | 0 (0) | 0 (0) | 0 |
| Frida | 111,098 | 97,111 (377) | 4,367 (54) | 2,830 |
| Phenol-Explorer | 7,486 | 7,486 (1,308*) | 7,486 (1,308*) | 5,285 |
| $FoodAtlas_{annotation}$ | 3,979 | 3,979 (1,385) | 3,979 (1,385) | 3,979 |
| $FoodAtlas_{prediction}$ | 225,902 | 225,902 (146,476) | 225,902 (146,476) | 225,902 |

*: All literature sources were curated from WoS according to the Phenol-Explorer website.

### 4 References

1. Hooton, F., Menichetti, G. & Barabási, A.-L. Exploring food contents in scientific literature with FoodMine. *Sci. Rep.* **10**, 16191 (2020).
2. Benjamini, Y. & Hochberg, Y. Controlling the False Discovery Rate: A Practical and Powerful Approach to Multiple Testing. *J. R. Stat. Soc. Ser. B Methodol.* **57**, 289–300 (1995).
3. Kim, S. *et al.* PubChem Protein, Gene, Pathway, and Taxonomy Data Collections: Bridging Biology and Chemistry through Target-Centric Views of PubChem Data. *J. Mol. Biol.* **434**, 167514 (2022).
4. KEGG: Kyoto Encyclopedia of Genes and Genomes | Nucleic Acids Research | Oxford Academic. https://academic.oup.com/nar/article/28/1/27/2384332.
5. UniProt: the Universal Protein knowledgebase | Nucleic Acids Research | Oxford Academic. https://academic.oup.com/nar/article/32/suppl_1/D115/2505378.
6. BLAST: Basic Local Alignment Search Tool. https://blast.ncbi.nlm.nih.gov/Blast.cgi.
7. Senanayake, U. M., Lee, T. H. & Wills, R. B. H. Volatile constituents of cinnamon (Cinnamomum zeylanicum) oils. *J. Agric. Food Chem.* **26**, 822–824 (1978).
8. Sommer, K., Hillinger, M., Eigenmann, A. & Vetter, W. Characterization of various isomeric photoproducts of ergosterol and vitamin D2 generated by UV irradiation. *Eur. Food Res. Technol.* **249**, 713–726 (2023).
9. Joradon, P. *et al.* Ergosterol Content and Antioxidant Activity of Lion's Mane Mushroom (Hericium erinaceus) and Its Induction to Vitamin D2 by UVC-Irradiation: in *Proceedings of the 8th International Conference on Agricultural and Biological*

*Sciences* 19–28 (SCITEPRESS - Science and Technology Publications, 2022). doi:10.5220/0011594600003430.

10. Dahlqvist, A. *et al.* Phospholipid:diacylglycerol acyltransferase: An enzyme that catalyzes the acyl-CoA-independent formation of triacylglycerol in yeast and plants. *Proc. Natl. Acad. Sci. U. S. A.* **97**, 6487–6492 (2000).

11. Shamsiev, A., Park, J., Olawuyi, I. F., Odey, G. & Lee, W. Optimization of ultrasonic-assisted extraction of polyphenols and antioxidants from cumin (Cuminum cyminum L.). *Korean J. Food Preserv.* **28**, 510–521 (2021).

12. Nagao, A., Maeda, M., Lim, B. P., Kobayashi, H. & Terao, J. Inhibition of β-carotene-15,15′-dioxygenase activity by dietary flavonoids. *J. Nutr. Biochem.* **11**, 348–355 (2000).

13. Shi, X. *et al.* Identification and investigation of a novel NADP+-dependent secoisolariciresinol dehydrogenase from Isatis indigotica. *Front. Plant Sci.* **13**, (2022).

14. Cui, H. *et al.* Comparative analysis of nuclear, chloroplast, and mitochondrial genomes of watermelon and melon provides evidence of gene transfer. *Sci. Rep.* **11**, 1595 (2021).

15. Wishart, D. S. *et al.* PathBank: a comprehensive pathway database for model organisms. *Nucleic Acids Res.* **48**, D470–D478 (2020).

16. Wishart, D. S. *et al.* HMDB 5.0: the Human Metabolome Database for 2022. *Nucleic Acids Res.* **50**, D622–D631 (2022).

17. McKillop, K., Harnly, J., Pehrsson, P., Fukagawa, N. & Finley, J. FoodData Central, USDA's Updated Approach to Food Composition Data Systems. *Curr. Dev. Nutr.* **5**, 596 (2021).

18. Neveu, V. *et al.* Phenol-Explorer: an online comprehensive database on polyphenol contents in foods. *Database* **2010**, bap024 (2010).

19. Rothwell, J. A. *et al.* Phenol-Explorer 2.0: a major update of the Phenol-Explorer database integrating data on polyphenol metabolism and pharmacokinetics in humans and experimental animals. *Database* **2012**, bas031 (2012).

20. Rothwell, J. A. *et al.* Phenol-Explorer 3.0: a major update of the Phenol-Explorer database to incorporate data on the effects of food processing on polyphenol content. *Database* **2013**, bat070 (2013).

21. Lee, J. *et al.* BioBERT: a pre-trained biomedical language representation model for biomedical text mining. *Bioinformatics* btz682 (2019) doi:10.1093/bioinformatics/btz682.

22. Wolf, T. *et al.* Transformers: State-of-the-Art Natural Language Processing. in *Proceedings of the 2020 Conference on Empirical Methods in Natural Language Processing: System Demonstrations* 38–45 (Association for Computational Linguistics, 2020). doi:10.18653/v1/2020.emnlp-demos.6.

23. Paszke, A. *et al.* PyTorch: An Imperative Style, High-Performance Deep Learning Library. in *Advances in Neural Information Processing Systems* vol. 32 (Curran Associates, Inc., 2019).

24. Loshchilov, I. & Hutter, F. Decoupled Weight Decay Regularization. Preprint at https://doi.org/10.48550/arXiv.1711.05101 (2019).

25. Settles, B. *Active Learning Literature Survey*. https://minds.wisconsin.edu/handle/1793/60660 (2009).

26. Bordes, A., Usunier, N., Garcia-Duran, A., Weston, J. & Yakhnenko, O. Translating Embeddings for Modeling Multi-relational Data. in *Advances in Neural Information Processing Systems* vol. 26 (Curran Associates, Inc., 2013).

27. Dong, X. *et al.* Knowledge vault: a web-scale approach to probabilistic knowledge fusion. in *Proceedings of the 20th ACM SIGKDD international conference on Knowledge discovery and data mining* 601–610 (Association for Computing Machinery, 2014). doi:10.1145/2623330.2623623.

28. Yang, B., Yih, W., He, X., Gao, J. & Deng, L. Embedding Entities and Relations for Learning and Inference in Knowledge Bases. Preprint at https://doi.org/10.48550/arXiv.1412.6575 (2015).

29. Ji, G., He, S., Xu, L., Liu, K. & Zhao, J. Knowledge Graph Embedding via Dynamic Mapping Matrix. in *Proceedings of the 53rd Annual Meeting of the Association for Computational Linguistics and the 7th International Joint Conference on Natural Language Processing (Volume 1: Long Papers)* 687–696 (Association for Computational Linguistics, 2015). doi:10.3115/v1/P15-1067.

30. Trouillon, T., Welbl, J., Riedel, S., Gaussier, E. & Bouchard, G. Complex Embeddings for Simple Link Prediction. in *Proceedings of The 33rd International Conference on Machine Learning* 2071–2080 (PMLR, 2016).

31. Sun, Z., Deng, Z.-H., Nie, J.-Y. & Tang, J. RotatE: Knowledge Graph Embedding by Relational Rotation in Complex Space. Preprint at http://arxiv.org/abs/1902.10197 (2019).

32. Ali, M. *et al.* PyKEEN 1.0: a Python library for training and evaluating knowledge graph embeddings. *J. Mach. Learn. Res.* **22**, 82:3723-82:3728 (2021).

33. OpenAI. ChatGPT. https://openai.com/blog/chatgpt.

34. Technical University of Denmark. Frida Food Data, Version 4.2. https://frida.fooddata.dk/?lang=en.

35. *Entrez Programming Utilities Help*. (National Center for Biotechnology Information (US), 2010).
